# Mitophagy enhancers against phosphorylated Tau-induced mitochondrial and synaptic toxicities in Alzheimer disease

**DOI:** 10.1101/2021.09.21.461119

**Authors:** Sudhir Kshirsagar, Neha Sawant, Hallie Morton, Arubala P. Reddy, P. Hemachandra Reddy

**Affiliations:** Department of Internal Medicine, Texas Tech University Health Sciences Center, Lubbock, TX 79430, USA; Nutritional Sciences Department, College of Human Sciences, Texas Tech University, 1301 Akron Ave, Lubbock TX 79409, USA; Department of Pharmacology and Neuroscience, Texas Tech University Health Sciences Center, Lubbock, TX 79430, USA; Department of Neurology, Texas Tech University Health Sciences Center, Lubbock, TX 79430, USA; Department of Public Health, Graduate School of Biomedical Sciences, Texas Tech University Health Sciences Center, Lubbock, TX 79430, USA; Department of Speech, Language, and Hearing Sciences, Texas Tech University Health Sciences Center, Lubbock, TX 79430, USA

**Keywords:** Mitophagy enhancers, Mitochondria, Synaptic activity, Urolithin A, Mitochondrial fragmentation

## Abstract

The purpose of our study is to determine the protective effects of mitophagy enhancers against phosphorylated tau (P-tau)-induced mitochondrial and synaptic toxicities in Alzheimer’s disease (AD). Mitochondrial abnormalities, including defective mitochondrial dynamics, biogenesis, axonal transport and impaired clearance of dead mitochondria are linked to P-tau in AD. Mitophagy enhancers are potential therapeutic candidates to clear dead mitochondria and improve synaptic and cognitive functions in AD. We recently optimized the doses of mitophagy enhancers urolithin A, actinonin, tomatidine, nicotinamide riboside in immortalized mouse primary hippocampal (HT22) neurons. In the current study, we treated mutant Tau expressed in HT22 (mTau-HT22) cells with mitophagy enhancers and assessed mRNA and protein levels of mitochondrial/synaptic genes, cell survival and mitochondrial respiration. We also assessed mitochondrial morphology in mTau-HT22 cells treated and untreated with mitophagy enhancers. Mutant Tau-HT22 cells showed increased fission, decreased fusion, synaptic & mitophagy genes, reduced cell survival and defective mitochondrial respiration. However, these events were reversed in mitophagy enhancers treated mTau-HT22 cells. Cell survival was increased, mRNA and protein levels of mitochondrial fusion, synaptic and mitophagy genes were increased, and mitochondrial fragmentation is reduced in mitophagy enhancers treated mTau-HT22 cells. Further, urolithin A showed strongest protective effects among all enhancers tested in AD. Our combination treatments of urolithin A + EGCG, addition to urolithin A and EGCG individual treatment revealed that combination treatments approach is even stronger than urolithin A treatment. Based on these findings, we cautiously propose that mitophagy enhancers are promising therapeutic drugs to treat mitophagy in patients with AD.

## Introduction

Alzheimer’s disease (AD) is a late-onset, progressive, mental illness, characterized by the progressive decline of multiple cognitive impairments with no cure ^1,2^. Currently, over 50 million people are affected globally by AD, of which more 5.8 million are living in United States Furthermore, this number is projected to increase to reach 7.1 million by the year 2025 and 13.8 million Americans by the year 2050 (Alzheimer’s Association, Facts and Figures 2020). AD is now the 6th leading cause of death in United States and is a major growing health concern world-wide.

Several years of intense research on postmortem AD brains and cell and mouse models of AD have revealed that multiple cellular changes are involved with the disease process, including deregulation of microRNAs, proliferation of astrocytes & synaptic loss/damage, increased production of free radicals & lipid peroxidation, increased mitochondrial fragmentation, in addition to increased accumulation of phosphorylated tau (P-tau) and amyloid beta (Aβ) ^3-16^. These changes were primarily observed in learning and memory regions, including hippocampus and cortex of the brain. Loss of synapses and synaptic damage are the best correlates of cognitive decline in patients diagnosed with AD and these tightly linked to P-tau & Aβ and mitochondria.

Tau is a major microtubule-associated protein that plays a large role in the outgrowth of neuronal processes and the development of neuronal polarity. Tau promotes microtubule assembly, stabilizes microtubules, promotes axonal transport, and affects the dynamics of microtubules in neurons ^17-19^. However, when tau becomes hyperphosphorylated, loses its microtubule binding ability, leading to impaired axonal transport and disrupt the cytoskeleton of neurons, causing defective synaptic transmission, dysfunction, and neuronal death ^20^. Hyperphosphorylated tau will form paired helical filaments which will then aggregate to form the neurofibrillary tangles, characteristic of AD ^20-22^. Studies with transgenic mouse models of AD suggests that Aβ toxicity is mediated by tau. It has also been shown that Aβ plays a role in triggering the hyperphosphorylation of tau, suggesting that generation of Aβ precedes the accumulation of P-tau ^20^. Oxidative stress from other sources such as decreased levels of insulin-like growth factor 1 have also been implicated in the formation of P-tau, leading to decreased cell viability. Loss of cytoskeletal integrity and subsequent neuronal death thus leads to the symptoms associated with AD ^20-22^. Recent reports demonstrated that Aβ-induced oxidative stress and other factors are critically involved in tau hyperphosphorylation in AD ^23-25^. Increasing evidence also suggests that tau is associated with mitochondria, particularly in disease state such as AD, leading to mitochondrial abnormalities in disease progression ^9^.

Mitochondrial abnormalities, including changes in mitochondrial DNA, mitochondrial gene expressions, mitochondrial ATP, mitochondrial enzymatic activities, mitochondria-induced increased free radicals, mitochondria-induced lipid peroxidation, reduced axonal transport, impaired biogenesis & dynamics and defective mitophagy are extensively reported in progression and pathogenesis of AD ^4-17, 26-37^. These changes are age-dependent, irrespective of type of AD - early-onset familial and late-onset sporadic. However, in familial AD, these changes occur much earlier than late-onset AD, because of the presence of genetic mutation(s) in amyloid beta precursor protein (APP) presenilin 1 (PS1) and presenilin 2 (PS2) alleles.

As stated above, current research suggests that mitochondrial damage is an important early cellular change in AD. Further, emerging evidence suggests that structural changes in mitochondria, including increased mitochondrial fragmentation and decreased mitochondrial fusion, are critical factors associated with mitochondrial dysfunction and cell death in aging and age-related diseases, including AD ^9,6^.

Mitochondrial dynamics is a delicate balance between division (fission) and fusion that maintains the shape and structure of mitochondria. In healthy cells, fission and fusion events are equally balanced, which maintains mitochondrial function ^9,15,38,39^. However, in AD and other neurological diseases such as HD, PD, ALS and others, mitochondrial dynamics is impaired, mostly with increased fragmentation and reduced fusion, leading to reduced PINK1 and Parkin levels, ultimately leading to defective clearance of dead mitochondria ^9,40^.

Mitochondrial biogenesis is the e synthesis of new mitochondria in the cell. There are four genes that are involved in mitochondrial biogenesis: PGC1α (PPAR – peroxisome proliferator-activated receptor)-γ coactivator-1α), NRF1 (nuclear respiratory factor 1), NRF2 (nuclear respiratory factor 2) and TFAM (transcription factor A, mitochondrial) ^41,42^. Mitochondrial biogenesis is affected by aging, age-related accumulation of mtDNA changes and toxins. In a diseased state, such as AD, mitochondrial biogenesis is defective, mainly because of the mutant protein(s) such as Aβ, p-tau association with mitochondria and increased free radical production and reduced mitochondrial function ^9^. Recent evidence from our lab and others indicates that mitophagy proteins PINK1 and Parkin are largely reduced in AD, primarily due to abnormal interactions between Aβ, p-tau with fission protein Drp1, leading to reduced clearance of dead mitochondria.

Mitophagy is the clearance of dead mitochondria by autophagy. Mitophagy is initiated by autophagosome. Autophagosomes deliver cytoplasmic components by lysosomes ^8,9,43^. The outer membrane of an autophagosome fuses with a lysosome to then form an autolysosome where the enveloped contents are subsequently degraded or cleared. In recent years, much progress has been made on this issue, and studies have suggested that several different organelles and potential membrane sources are the key to the initiation of this process. These include the plasma membrane, the Golgi apparatus, the ER and mitochondria. Many studies reported that autophagosomes are formed by the ER-mitochondria in mammalian cells.

Impaired mitochondrial dynamics, defective mitochondrial biogenesis and reduced mitophagy are reported in AD and other neurological diseases ^9, 44^. These events lead to increased proliferation of dead or dying mitochondria in neurons, particularly in hippocampal and cortical regions that are responsible for learning and memory in AD patients. Removal and/or clearance of these dead or dying mitochondria is important to maintain mitochondrial and synaptic functions in AD. Newly developed, pharmacological treatments and novel chemical modulators like nicotinamide riboside, tomatidine, actinonin, and urolithin A and even combination of mitophagy enhancers with amyloid beta and/or P-tau suppressors (such as green leaf extract EGCG) that might be used to promote the efficient removal of damaged mitochondria and restore the energetic status in AD neurons. Therapeutic drugs and natural supplements which targeting the mitophagy process could be a potential strategy to enhance the age-related disorders, which are characterized by mitophagy enhances when compared to defective controls. Defective mitophagy can be reduced through some natural supplementals, drugs to trigger mitochondrial events in a cell and enhance the clearance of dead and/or dying mitochondria.

We hypothesize that mitophagy can be enhanced by treating pharmacological enhancers such as nicotinamide riboside, tomatidine, actinonin, and urolithin A. To test our hypothesis, in the current study, we studied the protective effects of mitophagy enhancers against mutant Tau induced mitochondrial and synaptic toxicities in immortalized mouse hippocampal neurons (HT22). We transfected HT22 cells with mutant Tau cDNA and treated them with previously optimized dose for each mitophagy enhancers, nicotinamide riboside, tomatidine, actinonin, and urolithin A ^40^ and assessed mRNA and protein levels of mitochondrial dynamics, mitochondrial biogenesis, mitophagy/autophagy and synaptic genes. We also assessed cell survival and mitochondrial respiration, mitochondrial morphology in mTau-HT22 cells treated and untreated with mitophagy enhancers.

## Material and Methods

### Chemicals and Reagents

We got HT22 cells from David Schubert as a gift. Other cell culture supplements such as Dulbecco’s Modified Eagle Medium (DMEM), Minimum Essential Medium (MEM), penicillin/streptomycin, fetal bovine serum and Trypsin-EDTA were purchased from GIBCO (Gaithersburg, MD, USA).

### Mutant Tau cDNA constructs

Mutant Tau cDNA clone (P301L) has been purchased from Add gene - https://www.addgene.org and verified expression of mutations P301L and further sub-cloned into a mammalian expression vector. Western blot analysis was used to detect mutant tau protein expression to verify the expression of mutant Tau cDNA clone (P301L). Later, transfection of mutant Tau cDNA clone (P301L) into HT22 cells was done using lipofectamine 3000 for 24 hrs. Afterword’s, cells were treated with mitophagy enhancers (nicotinamide riboside, tomatidine, urolithin A and actinonin) (Sigma/Aldrich, CA) for 24hrs, then cells were harvested and pellet was collected to extract the RNA and proteins for further experiments.

### Tissue culture work

As per the standard cell culture, HT22 cells were grown for 3 days until they are 60-70% confluent. The medium used for growing these cells is 1:1 mixture of DMEM and OptiMEM with 10% FBS plus penicillin and streptomycin [Invitrogen, Carlsbad, CA, USA]. We have done 6 independent cell cultures and transfections with mutant Tau cDNA treatments for all experiments (HT22 cells, HT22 cells+mTau cDNA, HT22 cells+mTau cDNA+nicotinamide riboside, HT22 cells+mTau cDNA+tomatidine, HT22 cells+mTau cDNA+urolithin A and HT22 cells+mTau cDNA+Actinonin) (n=6) and later treated with mitophagy enhancers for 24 hrs.

### qRT-PCR analysis

Real time RT-PCR was carried out to check the mRNA expression of mitochondrial biogenesis, mitochondrial dynamics, mitophagy and synaptic genes. Total RNA was isolated from the HT22 cells using TriZol (Invitrogen, Carlsbad, CA) reagent. These cells were transfected with mutant tau cDNA for 24 hours and treated with mitophagy enhancers such as nicotinamide riboside, tomatidine, urolithin A and actinonin. To design the oligonucleotide primers, Primer Express Software (Applied Biosystems, Foster City, CA) was used. The primers were designed for the mitochondrial dynamic’s genes (Drp1, Fis1, Mfn1, Mfn2 and Opa1), mitochondrial biogenesis genes (PGC1α, NRF1, NRF2 and TFAM), mitophagy genes (PINK1), synaptic genes (synaptophysin) and housekeeping genes β-actin. SYBR-Green based quantitative real-time RT-PCR (ThermoFisher Scientific, Waltham, MA) was used for all the experiments.

Method includes DNAs treated total 5µg RNA as a starting material. Master mix of 1µl oligo dT, 19mM dNTPs (1µl), 5× first strand buffer (4 μl), 0.1 M DTT (2 μl) and 1 μl RNAseout was added to the starting material. In order to denature the RNA, Oligo dT, dNTPs, RNA, and other reagents were mixed first and then heated at 65°C for 5 minutes and then chilled on ice until the remaining components were added. Samples were incubated at 42°C for 2 minutes and added 1 μl of Superscript III (40 U/μl) to the mixture. Further, the samples were incubated at 42°C for 50 minutes and then samples were incubated at 70°C for 15 minutes in order to inactivate the reaction. Assay was done with triplicate samples using diluted cDNA of 100ng per 20 ul reaction, to run the sample QuantStudio3 (Applied Biosystems, Foster City, CA) was used. Temperature conditions used to run the PCR was; 50°C for 2 minutes and 95°C for 10 minutes, followed by 40 cycles of 95°C for 15 seconds and 60°C for 1 minute. The fluorescent spectra were documented at the elongation phase of each PCR cycle and a dissociation curve was created to distinguish non-specific amplicons. CT values were calculated by using Quant studio. The CT values and amplification plots was exported to Microsoft Excel worksheet for further analysis. As a housekeeping gene, β-actin was used and mRNA transcript levels were normalized against it for each dilution. As per CT method (Applied Biosystems, Foster City, CA), the relative quantification of gene of interest using housekeeping gene was calculated.

### Western blotting analysis

To perform western blotting analysis, HT22 cells were transfected with mTau cDNA and treated with mitophagy enhancers (nicotinamide riboside, tomatidine, urolithin A and actinonin) for 24hrs. Then cells were lysed in 50μl cold RIPA lysis buffer (Millipore Sigma Aldrich Corporation, 20–188) for 60 mins on ice (vortex every 15 min interval) and centrifuged at 12,000 g for 11 min. The supernatant was collected, and protein concentration was measured. 40μg proteins were loaded and separated by SDS-PAGE gels (10%) electrophoretically and transferred to polyvinylidene difluoride membrane (Bio-Rad Incorporation, 10026933). Membrane was blocked with 5% BSA for 60 min at room temperature on shaker. Primary antibody was added to the membranes after 3 washes with TBST solution and kept overnight at 4-degree temperature on shaker. Beta-actin was used as an internal control in order to outline the levels of mitochondrial biogenesis, dynamics, synaptic and mitophagy. Details of antibody dilutions are given in Table 3. Next day, membrane was washed 3 times with TBST and incubated with HRP (horseradish peroxidase)-labeled secondary antibodies for 1 hr at room temperature. To detect the proteins chemiluminescence reagents (ECL, Thermo scientific, WA317048) were used and the band exposures were kept within the linear range.

**Table 1.**
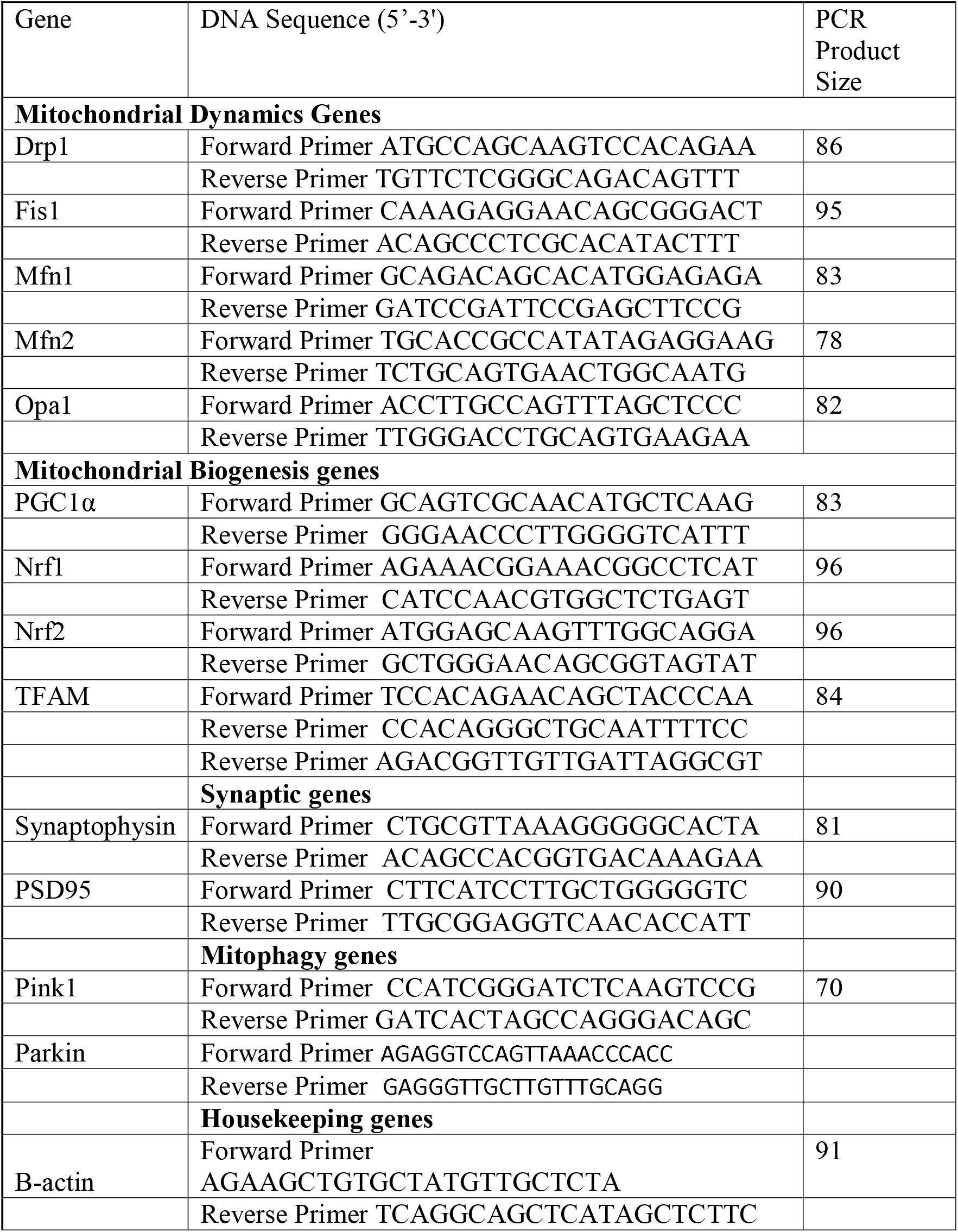
**Summary of qRT-PCR oligonucleotide primers used in measuring mRNA expressions in mitochondrial dynamics and mitochondrial biogenesis, synaptic and mitophagy genes in mitophagy enhancers treated and untreated mutant Tau HT22 cells**

**Table 2.** **Summary of antibody dilutions and conditions used in the immunoblotting analysis of mitochondrial dynamics, mitochondrial biogenesis, synaptic and mitophagy proteins in mitophagy enhancers treated and untreated mTau-HT22 cells and untransfected HT22 cells**

| **Marker Primary antibody—species and dilution** | **Purchased from company, city and state** | **Secondary antibody, dilution** | **Purchased from company, city and state** |
| --- | --- | --- | --- |
| Anti-Tau (phospho S202) antibody Rabbit monoclonal 1:500 | Abcam, Cambridge, MA | Donkey anti-rabbit HRP 1:10 000 | GE Healthcare Amersham, Piscataway, NJ |
| Drp1 Rabbit polyclonal 1:500 | Novus Biological, Littleton, CO | Donkey anti-rabbit HRP 1:10 000 | GE Healthcare Amersham, Piscataway, NJ |
| Fis1 Rabbit polyclonal 1:500 | Protein Tech Group, Inc, Chicago, IL | Donkey anti-rabbit HRP 1:10 000 | GE Healthcare Amersham, Piscataway, NJ |
| Mfn1 Rabbit polyclonal 1:400 | Abcam, Cambridge, MA | Donkey anti-rabbit HRP 1:10 000 | GE Healthcare Amersham, Piscataway, NJ |
| Mfn2 Rabbit polyclonal 1:400 | Abcam, Cambridge, MA | Donkey anti-rabbit HRP 1:10 000 | GE Healthcare Amersham, Piscataway, NJ |
| OPA1 Rabbit polyclonal 1:500 | Novus Biological, Littleton, CO | Donkey anti-rabbit HRP 1:10 000 | GE Healthcare Amersham, Piscataway, NJ |
| SYN Rabbit monoclonal 1:400 | Abcam, Cambridge, MA | Donkey anti-rabbit HRP 1:10 000 | GE Healthcare Amersham, Piscataway, NJ |
| PGC1a Rabbit polyclonal 1:500 | Novus Biological, Littleton, CO | Donkey anti-rabbit HRP 1:10 000 | GE Healthcare Amersham, Piscataway, NJ |
| NRF1 Rabbit polyclonal 1:300 | Novus Biological, Littleton, CO | Donkey anti-rabbit HRP 1:10 000 | GE Healthcare Amersham, Piscataway, NJ |
| NRF2 Rabbit polyclonal 1:300 | Novus Biological, Littleton, CO | Donkey anti-rabbit HRP 1:10 000 | GE Healthcare Amersham, Piscataway, NJ |
| TFAM Rabbit polyclonal 1:300 | Novus Biological, Littleton, CO | Donkey anti-rabbit HRP 1:10 000 | GE Healthcare Amersham, Piscataway, NJ |
| PINK1 Rabbit polyclonal 1:500 | Novus Biological, Littleton, CO | Donkey anti-rabbit HRP 1:10 000 | GE Healthcare Amersham, Piscataway, NJ |
| Parkin Mouse polyclonal 1:500 | Novus Biological, Littleton, CO | Sheep anti-mouse HRP 1:10 000 | GE Healthcare Amersham, |
|  |  |  | Piscataway, NJ |

**Table 3.**
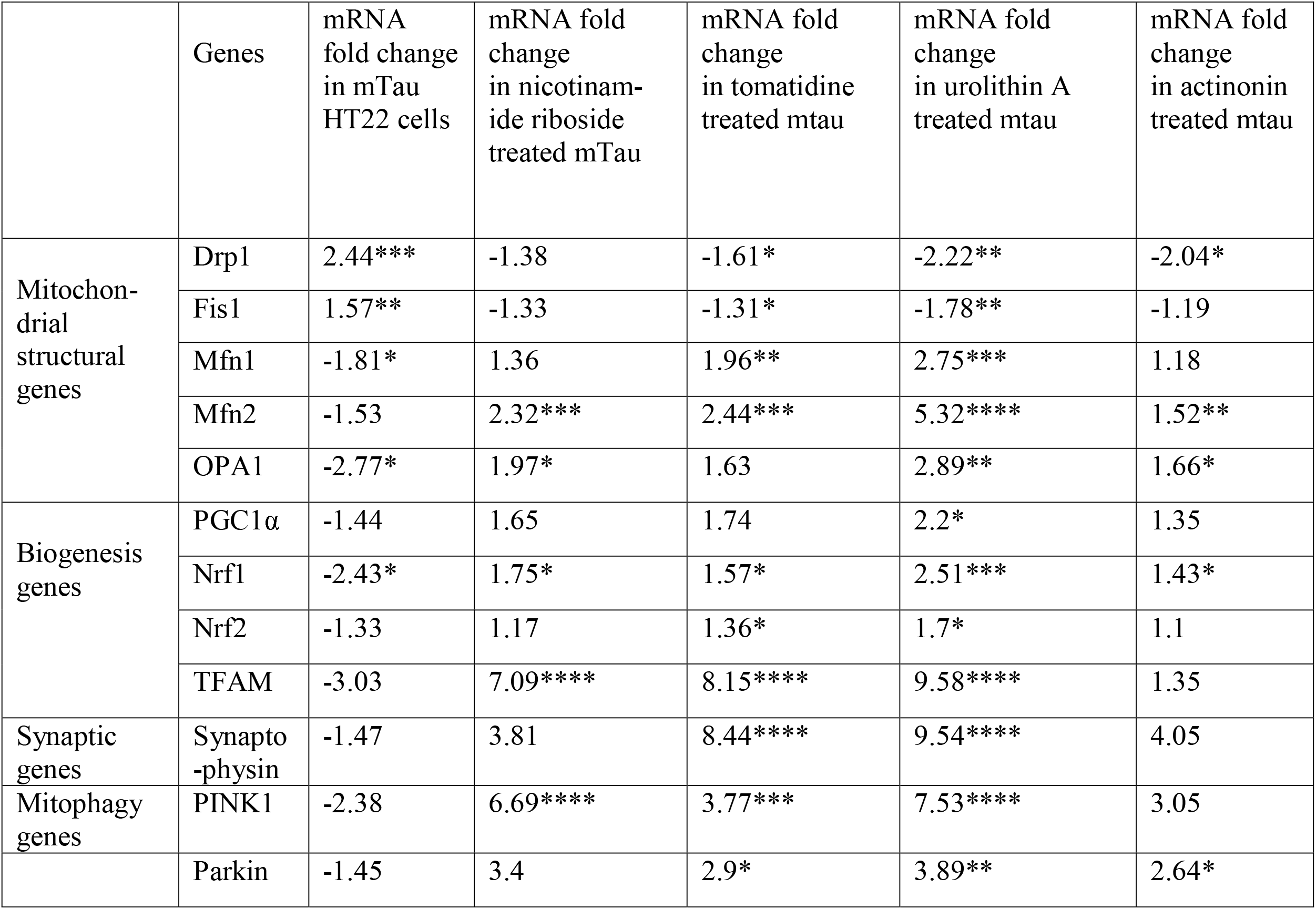
**Summary of mRNA fold changes in mutant Tau-HT22 cells relative control HT22 cells and mitophagy enhancers treated in mutant Tau-HT22 treated cells relative to mutant Tau-HT22 untreated cells**

### Cell survival/apoptotic assay

The cell survival assay was done as per user’s manual; this assay was done to check the cell apoptosis by using Cellometer K2 CBA Image Cytometry System (Nexcelom Bioscience LLC, Lawrence, MA). For the procedure, Annexin V-FITC and Propidium Iodide (PI) staining solution were used as fluorophore to detect the apoptotic and necrotic cells. HT22 cells were seeded and grown for overnight; cells were then harvested using trypsin and collected the pellet after centrifugation. Using Newbouwer chamber, cells were counted and 100,000 to 150,000 cells were collected and re-suspended in Annexin V binding buffer (40 µl). Next, Annexin V – FITC reagent (green) and PI (red) each 5 µl was added to the binding buffer containing cells. Then the mixture was gradually mixed by pipetting up and down for about 10 times and incubated at room temperature, in dark for 15 minutes. Mixture was then washed with 1XPBS (250 µl) by 3 minutes centrifugation, pellet was resuspended in 50 µl of Annexin V binding buffer and measured the cell apoptosis.

### Mitochondrial respiration using Seahorse XFe96 Extracellular Flux Analyzer

Mitochondrial respiration was measured by using advanced seahorse technique. For that HT22 cells were seeded in a petri dish and grown overnight, next day, cells were transfected with mTau plasmid for 24 hours. Cells were then trypsinized, counted and treated with mitophagy enhancers and plated 10,000 cells in 80 µL growth medium (DMEM medium supplemented with 10% fetal bovine serum, 1% penicillin and streptomycin) each well, except four background correction wells (A1, A12, H1, and H12), which should be blanked with 80 µL of growth medium. Then the plate was kept in cell culture hood to rest at 20°C–25°C for 1 hour, that way cells gets allocated uniformly and decrease edge effects for cells. Cell were then incubated in a cell culture incubator for overnight. The utility plate and sensory cartridge were detached and placed the sensory cartridge upside down on the bench side to the utility plate. Seahorse XF Calibrant (200 µL) was added to each well of the utility plate, then lower the sensory cartridge back onto the utility plate softly and avoided making air bubbles. The sensory cartridge was then kept in a non-CO_2_ and 37°C incubator for overnight. Next morning, the medium from XF96 cell culture microplate was removed leaving 20 µL of the media in the plate. Then added 180 µL of freshly prepared assay medium (1 mL 200 mM glutamine, 1 mL 100mM pyruvate solution, and 0.1 g D-glucose in 98 mL XF base medium), and incubated the XF96 cell culture microplate for 37°C in non-CO_2_ incubator for 1 hour. In between, the stock solutions of oligomycin, FCCP and rotenone/antimycin A were diluted as per the protocol and loaded 20 µL of 1.5 µM oligomycin in port A, 22 µL of 1 µM FCCP in port B and 25 µL of 0.5 µM rotenone/antimycin A in port C of the hydrated sensory cartridge. Later the utility plate was kept in the seahorse machine and hydrated sensory cartridge for calibration. After the calibration is done, the utility plate was removed and exchanged the XF96 cell culture microplate on the tray with the accurate direction as labeled on corner of the plate then loaded the tray. The total time of OCR measurements is 1 hour and 24 min. After the measurements are completed, the results are automatically formed, analyzed by the wave software and data was transferred to excel or prism file. Data shown are mean ± standard error of the mean from six to eight wells.

### Transmission electron microscopy

Transmission electron microscopy technique was used to measure the number and length of mitochondria in all control and experimental group of cells (untreated HT22, HT22 cells treated with mitophagy enhancers and cells transfected with mTau). To prepare the samples, first cells were fixed in 100 μm sodium cacodylate (pH 7.2), 2.5% glutaraldehyde, 1.6% paraformaldehyde, 0.064% picric acid and 0.1% ruthenium red. Cells were then gently washed and post-fixed in 1% osmium tetroxide plus 08% potassium ferricyanide, in 100 mm sodium cacodylate (pH 7.2) for 1 hour. The HT22 cells were then thoroughly rinsed in the water, dehydrated, infiltrated overnight in 1:1 acetone: Epon 812 and infiltrated for 1 hour with 100% Epon 812 resin and finally embedded in the resin. Once the polymerization is done, 60–80 nm thin sections were made on a Reichert ultramicrotome. Sections were stained for 5 min in lead citrate, rinsed and post-stained for 30 min in uranyl acetate. Later sections were again rinsed and dried. Electron microscopy was done at 60 kV on a Morgagni TEM Philips equipped with a CCD, and images were collected at magnifications of ×1000–37 000. The numbers of mitochondria and mitochondrial length were counted in all groups of cells. Statistical significance was determined using one-way analysis of variance.

## Results

### mRNA levels of mitochondrial dynamics and mitochondrial biogenesis genes

Messenger RNA (mRNA) levels of mitochondrial dynamics, mitochondrial biogenesis, mitophagy and synaptic genes were evaluated, in order to see the protective effect of mutant Tau toxicity and protective effects of mitophagy enhancers. For that, we used different groups of cells: 1) HT22 cells (control), 2) HT22 + mutant Tau cDNA transfected, 3) HT22 + mutant Tau cDNA transfected, and nicotinamide riboside treated, 4) HT22 + mutant Tau cDNA transfected and tomatidine treated, 5) HT22 + mutant Tau cDNA transfected and treated with urolithin A and 6) HT22 + Mutant Tau cDNA transfected and treated with actinonin (Figure 1). Using Sybr-Green chemistry-based quantitative real time RT-PCR, mRNA levels were measured. Those includes mitochondrial dynamic genes (fission Drp1 & Fis1 and fusion Mfn1, Mfn2 and Opa1), mitochondrial biogenesis genes (PGC1α, Nrf1, Nrf2 and TFAM), mitophagy (PINK1 and Parkin) and synaptic genes (synaptophysin).

**Figure 1:**
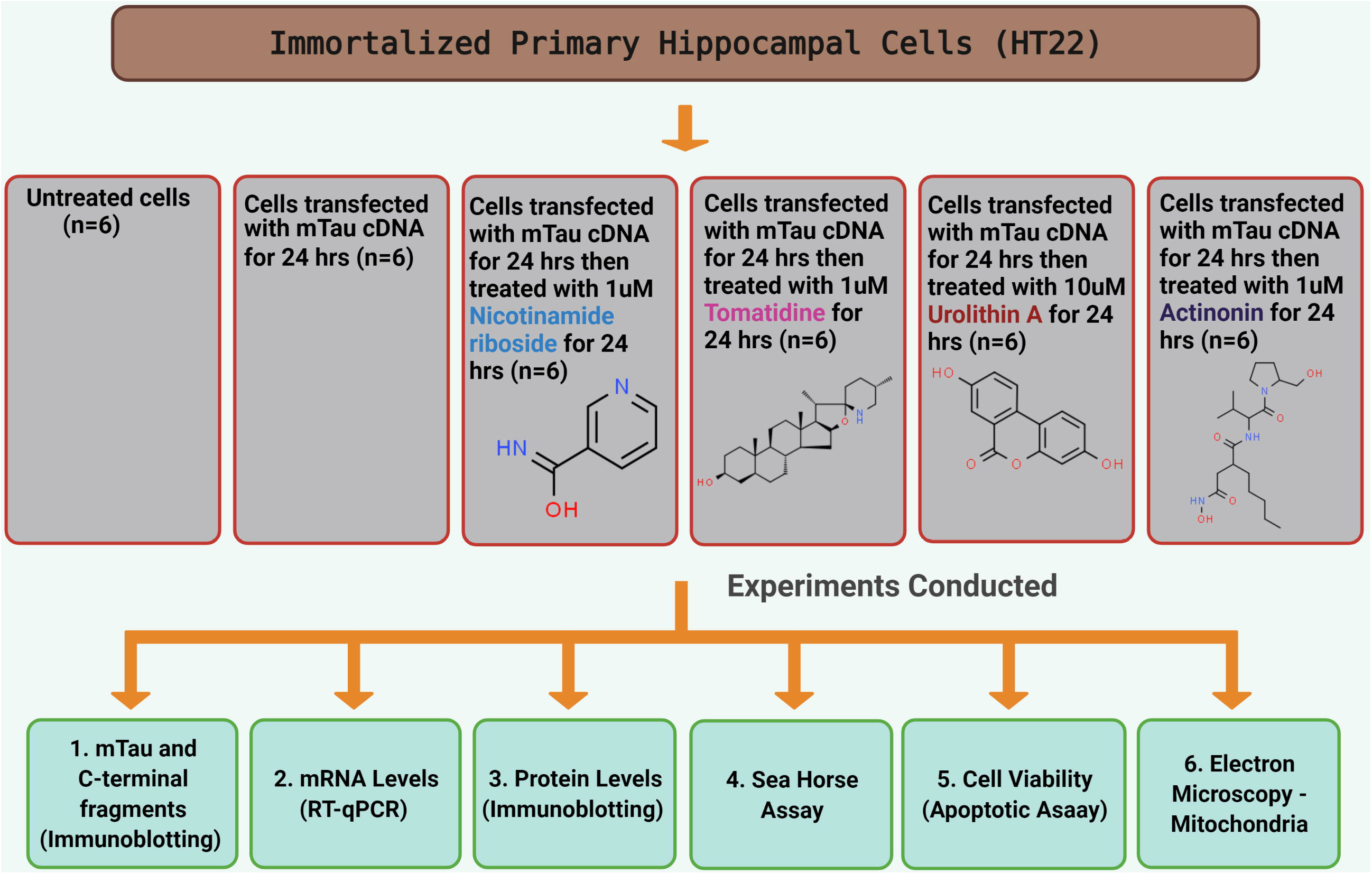
Flow chart of cells used in the present study for mitophagy enhancers (nicotinamide riboside, tomatidine, urolithin A and actinonin) treatment and experiment conducted.

#### Mitochondrial Dynamics

mRNA levels of mitochondrial fission genes were significantly increased (Drp1 by 2.4-fold and Fis1 by 1.5-fold) in the mutant Tau transfected HT22 cells (referred as mTau-HT22 from here on) as compared to HT22 cells. However, mRNA expression levels of mitochondrial fusion genes were significantly decreased (Mfn1 by 1.8-fold, Mfn2 by-1.5 fold and Opa1 by 2.7-fold) in mTau-HT22 cells compared to HT22 cells (Table 3). This observation specifies the presence of abnormal mitochondrial dynamics in mTau-HT22 cells.

Conversely, nicotinamide riboside treated mTau-HT22 cells showed reduced fission genes (Drp1 by 1.3-fold, Fis1 by 1.3-fold) and increased fusion genes (Mfn1 by 1.36-fold, Mfn2 by 2.3-fold, and Opa1 by 1.9-fold) (Table 3). These interpretations show that nicotinamide riboside reduces fission and increases fusion activity in mtau-HT22 cells.

Other mitophagy enhancers tomatidine, actinonin and urolithin A showed a similar findings (Table 3). However, urolithin A treated cells showed strong fold change differences, reduced fission genes expression (Drp1 by 2.2-fold, Fis1 by 1.7-fold) and increased fusion genes expressions (Mfn1 by 2.7-fold, Mfn2 by 5.3-fold, and Opa1 by 2.8-fold) (Table 3). These results indicate that urolithin A has strong mitophagy activity among all tested drugs in our study.

#### Mitochondrial biogenesis

In comparison with HT22 cells, mTau-HT22 cells exhibit significantly decreased mRNA levels of mitochondrial biogenesis genes (PGC1a by 1.4-fold; Nrf1 by 2.4; Nrf2 by 1.3-fold and TFAM by 3.0-fold) (Table3). On the other hand, in nicotinamide riboside treated mTau-HT22 cells, mRNA levels of mitochondrial biogenesis were increased (PGC1a by 1.6-fold; Nrf1 by 1.7-fold; Nrf2 by 1.1 and TFAM by 7.0-fold) representing that enhancers tomatidine, actinonin and urolithin A treated mTau-HT22 cells showed a similar pattern of mRNA expression levels. Urolithin A treated cells showed highly increased levels of all biogenesis genes (PGC1a by 2.2-fold, Nrf1 by 2.5-fold, Nrf2 by 1.7-fold and TFAM by 9.5-fold) (Table 3).

#### Mitophagy

As presented in Table 3, mRNA levels of mitophagy genes were reduced PINK1 by 2.3-fold and Parkin by 1.45-fold relative to control HT22 cells. On the other hand, mitophagy enhancers treated cells showed increased PINK1 levels; nicotinamide riboside by 6.6-fold, tomatidine by 3.7-fold, actinonin by 3.0-fold and urolithin A by 7.5-fold; Parkin levels nicotinamide riboside by 3.4-fold, tomatidine by 2.9-fold, actinonin by 2.64-fold and urolithin A by 3.8-fold. These findings show that mitophagy enhancers improved mitophagy activity in AD cells with a strong activity of urolithin A.

#### Synaptic

In comparison with HT22 cells, mTau-HT22 cells showed reduced synaptophysin mRNA levels by 1.4-fold (Table 3). However, synaptophysin levels were increased in mitophagy enhancers treated mTau-HT22 cells relative to mitophagy enhancers untreated mTau-HT22 cells (nicotinamide riboside by 3.8-fold, tomatidine by 8.4-fold, actinonin by 4.0-fold and urolithin A by 9.5-fold). These results shows that mitophagy enhancers increases synaptic activity in mTau-HT22 cells (Table 3).

### Immunoblotting Analysis: mitophagy enhancers reduce full-length mutant Tau

In order to understand the impact of mitophagy enhancers on mutant Tau, we studied mutant Tau-HT22 cells that were treated with mitophagy enhancers. As shown in Figure 2, a full-length 70-kDa pTau protein was found in the mutant tau cDNA transfected HT22 cells. The quantitative densitometry analysis of the full-length mutant Tau in transfected cells shows a bsignificant decrease in the mitophagy enhancers treated cells; for example, nicotinamide riboside (P=0.002), urolithin A (P=0.003), tomatidine (P=0.005) and actinonin (P=0.002). These observations suggests that mitophagy enhancers reduce mutant full-length mutant Tau.

**Figure 2:**
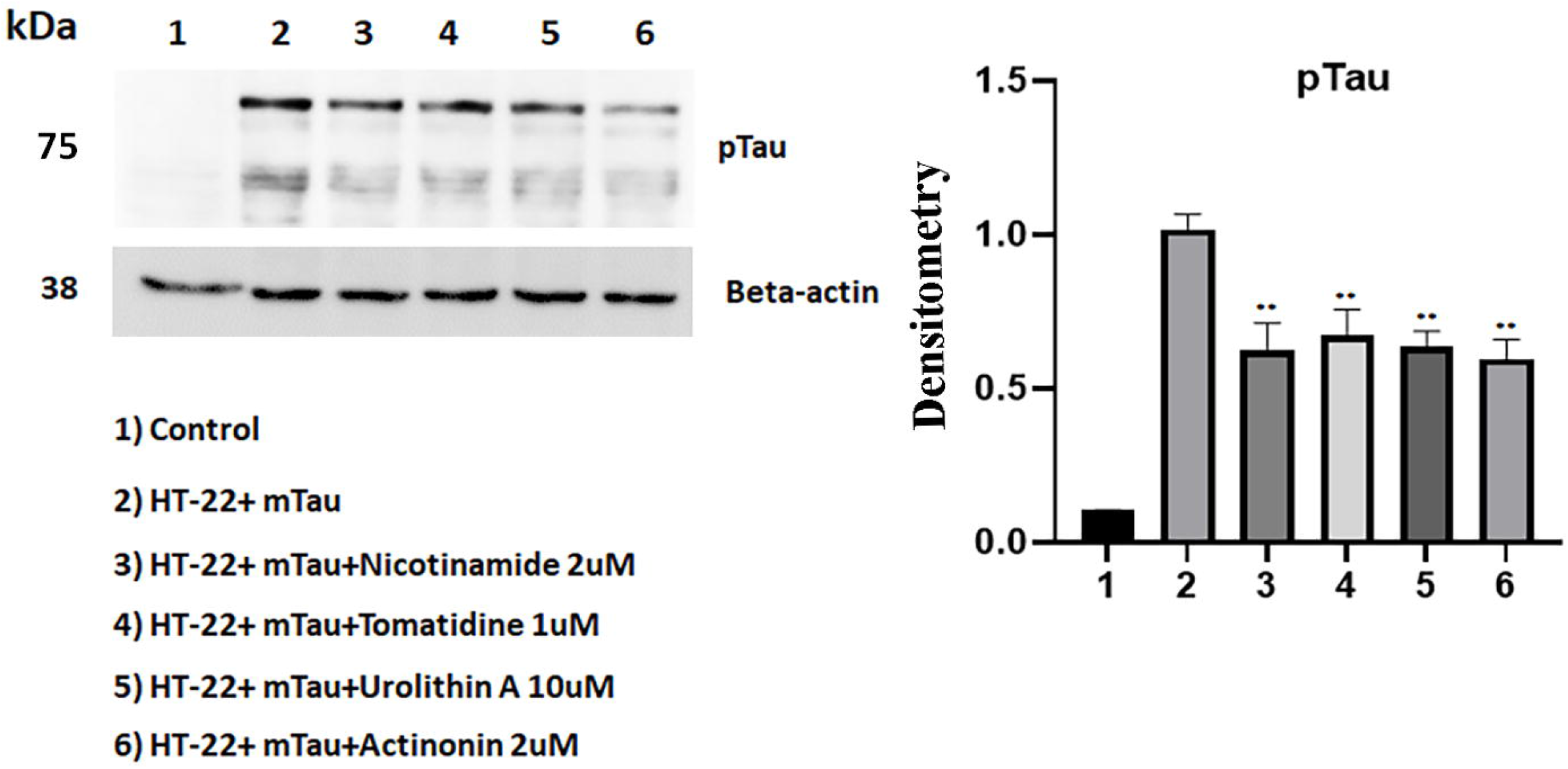
Immunoblotting of mTau-HT22 cells with pTau antibody. **(A)** A full-length 70 kDa mTau protein was found in the transfected cells. **(B)** Quantitative densitometry analysis of the full length mTau.

### Mitophagy enhancers increases/maintains mitochondrial dynamics

We performed immunoblotting analysis in order to determine the toxic effects of mutant tau against mitochondrial dynamics. Protein expressions of fission (Drp1 and Fis1, and fusion Mfn1, Mfn2 and Opa1) in mutant Tau-HT22 cells and control HT22 cells are shown in Figures 3. As compared to HT22 cells, mutant Tau-HT cells showed increased fission (Drp1 – P=0.01, Fis1 – P=0.005) and reduced fusion proteins expressions (Mfn2 – P=0.002 and Opa1 P=0.01).

**Figure 3:**
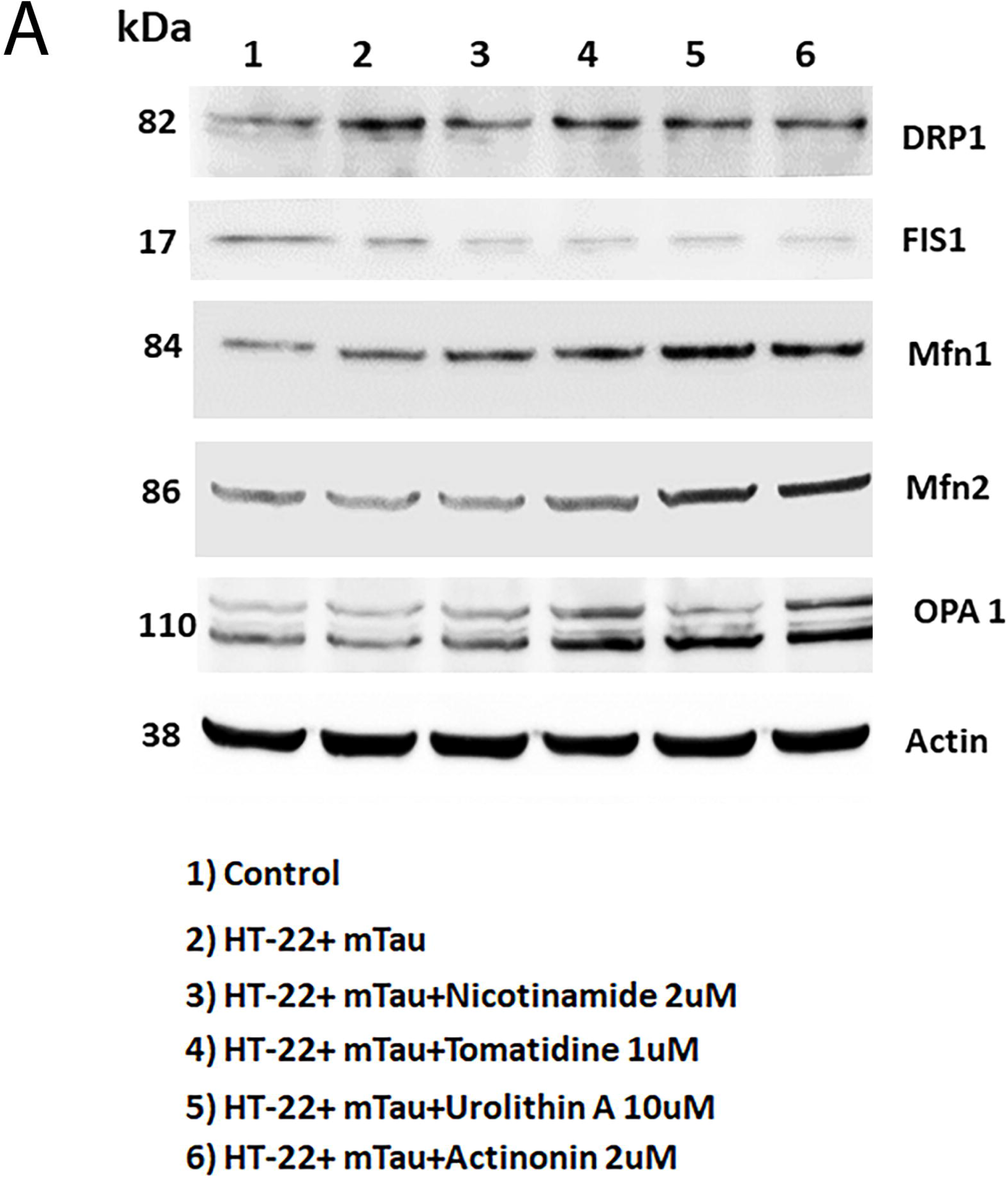
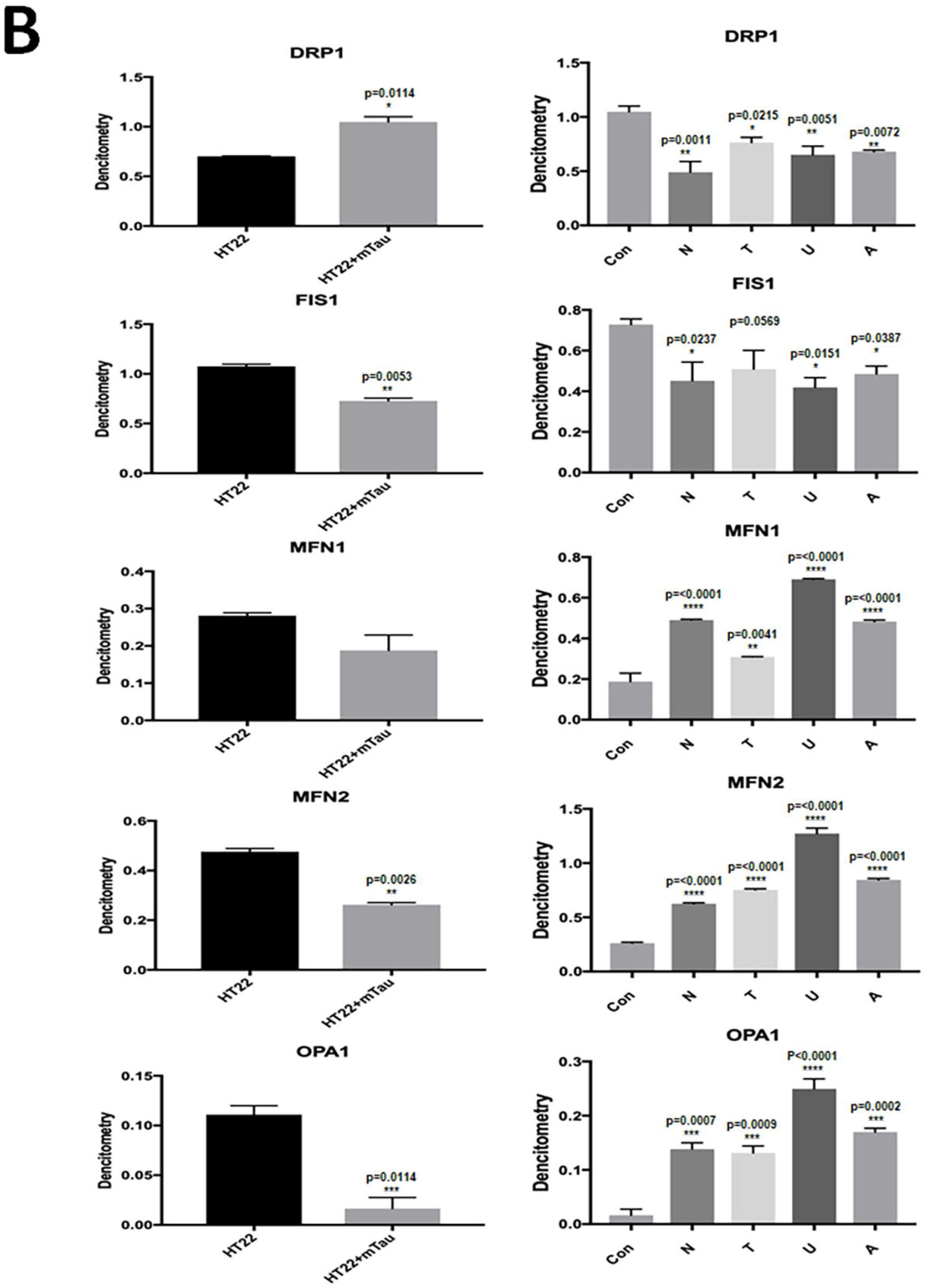
Western blot analysis of mitochondrial dynamics proteins. **(A)** Representative immunoblots for control and mTau-HT22 cells with or without mitophagy enhancers. **(B)** Quantitative densitometry analysis for mitochondrial dynamics proteins – significantly increased levels of fission proteins Drp1 and Fis1 were observed in cells transfected with mutant mTau. Fusion proteins Mfn1, Mfn2 and Opa1 were significantly decreased. On the other hand, mitophagy enhancers treated mutant Tau showed reduced levels of fission proteins and increased levels of fusion proteins were observed.

In contrast, decreased levels of fission proteins and increased levels of fusion proteins were found in mitophagy enhancers treated HT22 cells compared to untreated cells (Figure 3A 4 3B). P values of these proteins are; Drp1 – nicotinamide riboside P=0.001, tomatidine P=0.02, urolithin A P=0.005, actinonin P=0.007; Fis1-nicotinamide riboside P=0.02, tomatidine P=0.05, urolithin A P=0.01, actinonin P=0.03). Increased levels of fusion proteins are observed; Mfn1 - 0.0001 tomatidine P=0.004, urolithin A P=0.0001, actinonin P=0.0001, Mfn2 nicotinamide P=0.0001, tomatidine P=0.0001, urolithin A P=0.0001 & Opa1-urolithin A P=0.0001) (Figure 3B0. Importantly, urolithin A showed the highest protective effect among all mitophagy enhancers.

### Mitophagy enhancers increases mitochondrial biogenesis proteins

In order to determine the toxic effects of mutant tau against mitochondrial biogenesis and to see the protective effects of mitophagy enhancers against the mutant tau induced mitochondrial biogenesis, we performed immunoblotting analysis. As presented in Figure 4A, expression levels of mitochondrial biogenesis proteins were decreased in mTau-HT22 cells (PGC1a - P=0.005, NRF1 - P=0.0004, NRF2 - P=0.01 & TFAM – P=0.003) (Figure 4B).

**Figure 4:**
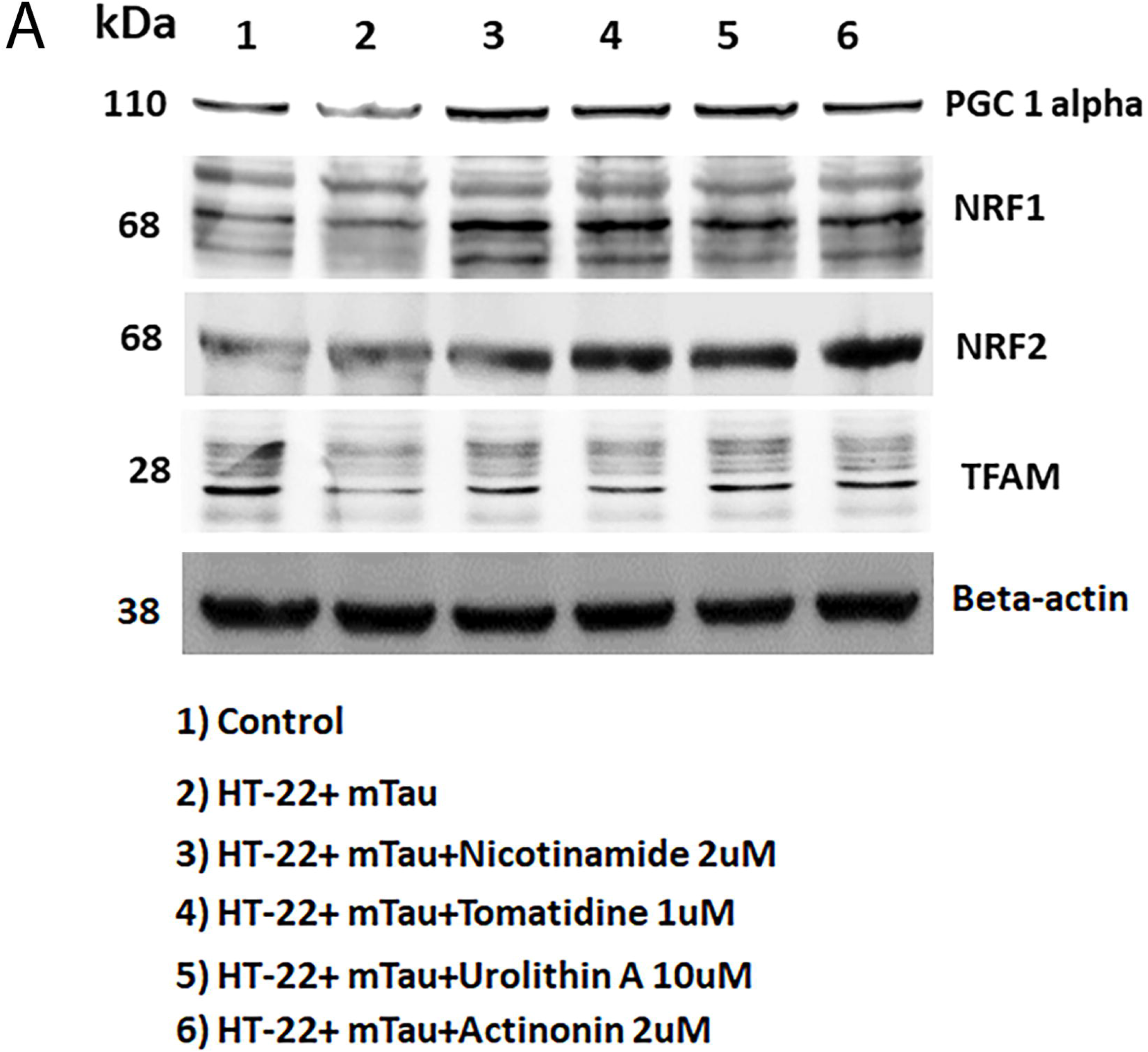
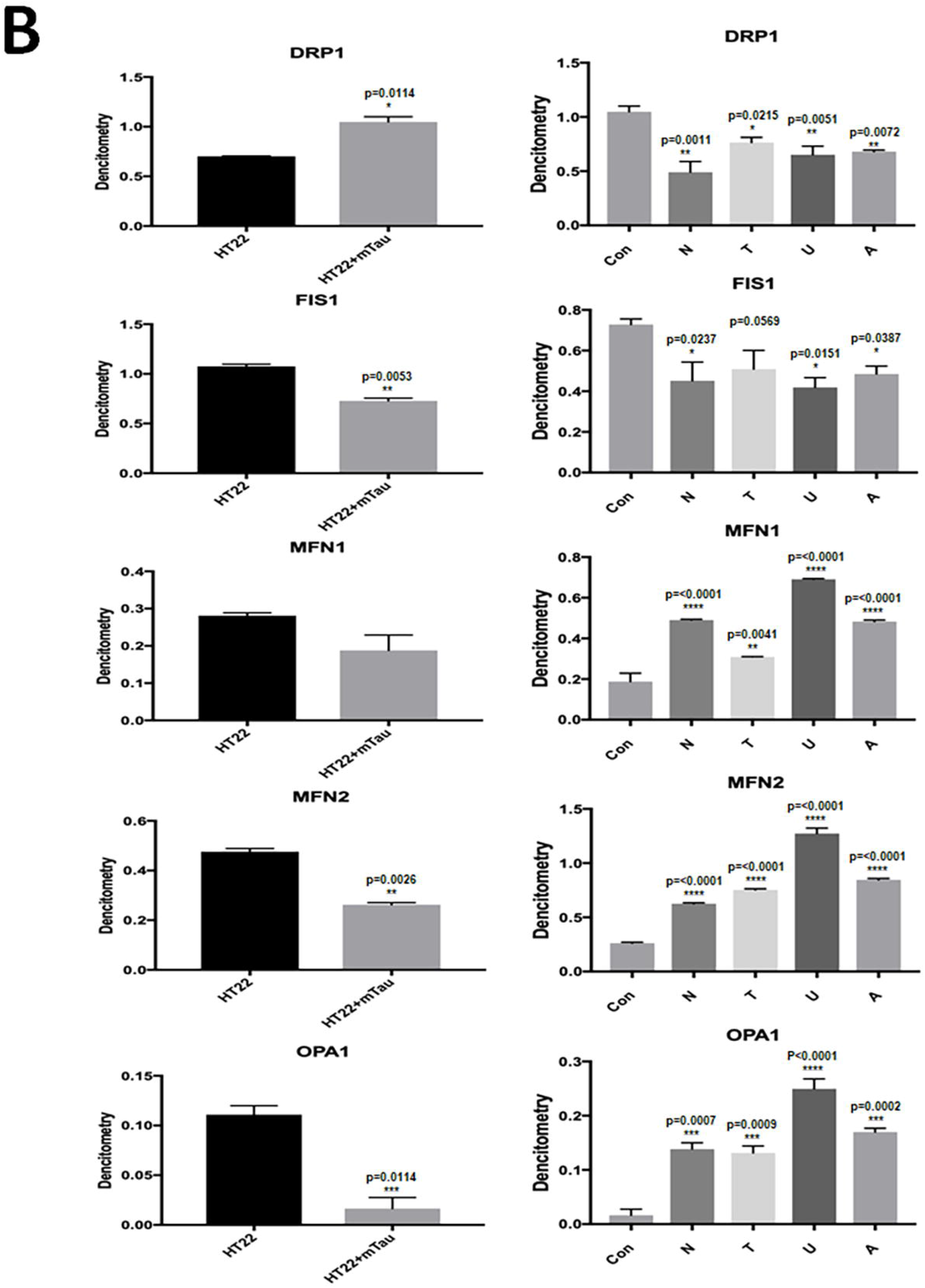
Western blot analysis of mitochondrial biogenesis proteins in HT22 cells and mutant Tau cDNA transfected and treated with mitophagy enhancers for 24 hours. **(A)** Representative immunoblots for control HT22 and mTau-HT22 cells with or without mitophagy enhancers treatment. **(B)** Quantitative densitometry analysis showed significant reduction in the levels of PGC1a, NRF1, NRF2 and TFAM upon mTau cDNA transfection. But levels of all mitochondrial biogenesis proteins increased with mitophagy enhancers treatment.

On the other hand, expression levels of mitochondrial biogenesis proteins were increased in mitophagy enhancers (nicotinamide riboside, tomatidine, urolithin A and actinonin) treated mTau-HT22 cells compared to mitophagy enhancers untreated mTau-HT22cells. P values for those proteins are; PGC1a - nicotinamide riboside P=0.0001; tomatidine P=0.0001; urolithin A P=0.0001 actinonin P=0.0001; NRF1 - nicotinamide riboside P=0.0001, tomatidine P=0.0001, urolithin A P=0.0001, actinonin P=0.0001; NRF2 - tomatidine P=0.04, urolithin A P=0.005, actinonin P=0.06 & TFAM – nicotinamide riboside P=0.0001, tomatidine P=0.0001, urolithin A P=0.0001, Actinonin P=0.0002.

### Mitophagy enhancers enhances mitophagy proteins

The expression of mitophagy proteins was detected in order to study the protective role of mitophagy enhancers (nicotinamide riboside, tomatidine, urolithin A and actinonin) against tau-induced PINK1 and parkin. Mutant Tau-HT22 cells showed reduced PINK1 P=0.03 and parkin protein expressions compared to control HT22 cells. Conversely, PINK1 and Parkin expression was increased in mutant Tau-HT22 cells treated with mitophagy enhancers, PINK1 nicotinamide riboside P=0.001, tomatidine P=0.0004, urolithin A P=0.0002 and actinonin P=0.01, Parkin nicotinamide riboside P=0.0001, tomatidine P=0.0001, urolithin A P=0.0001 and actinonin P=0.001 (Figure 5A and 5B). These results suggest that mitophagy enhancers increases expression of PINK1 and parkin proteins; most importantly, urolithin A treated cells showed highest expressions compared to other mitophagy enhancers.

**Figure 5:**
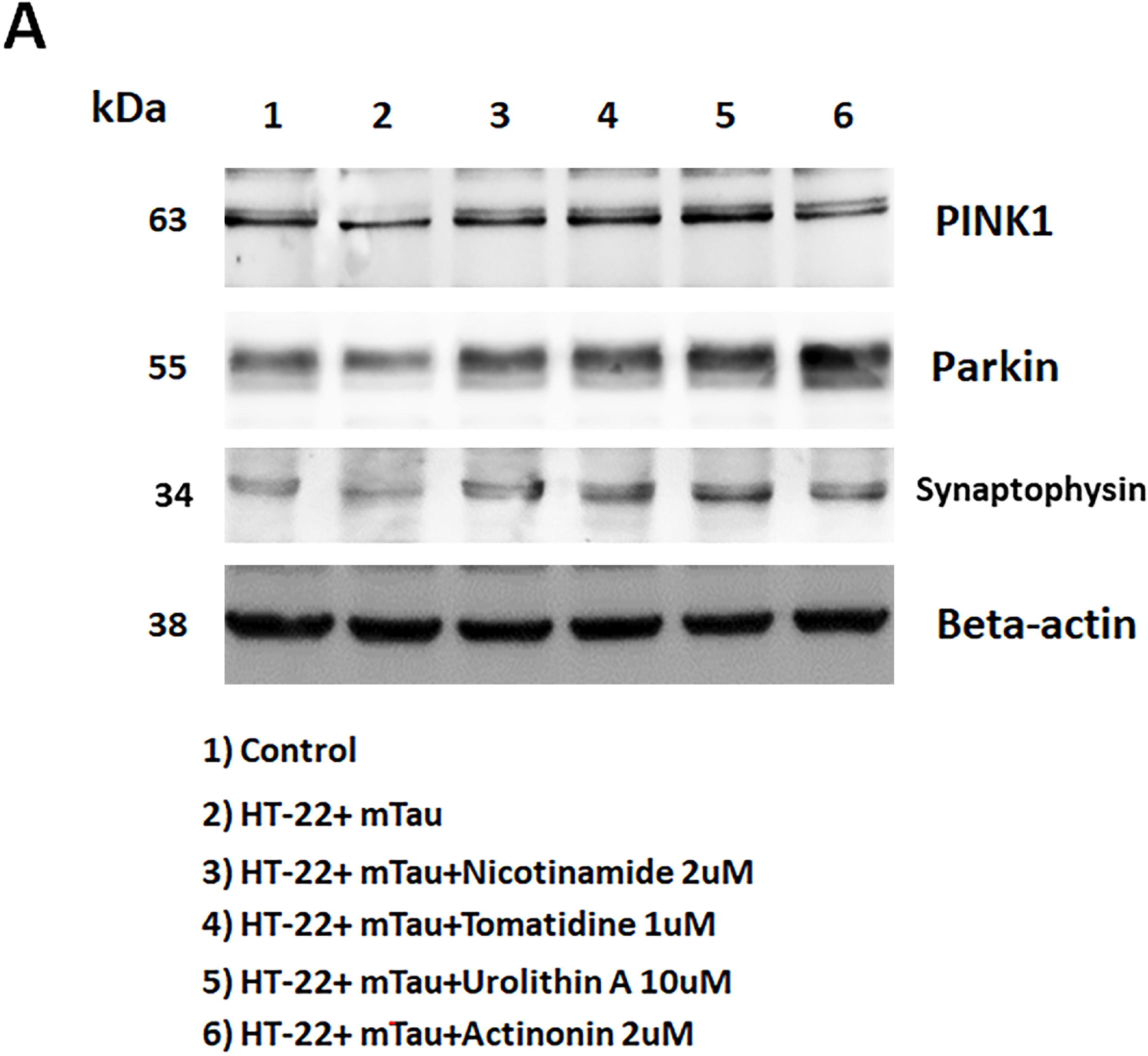
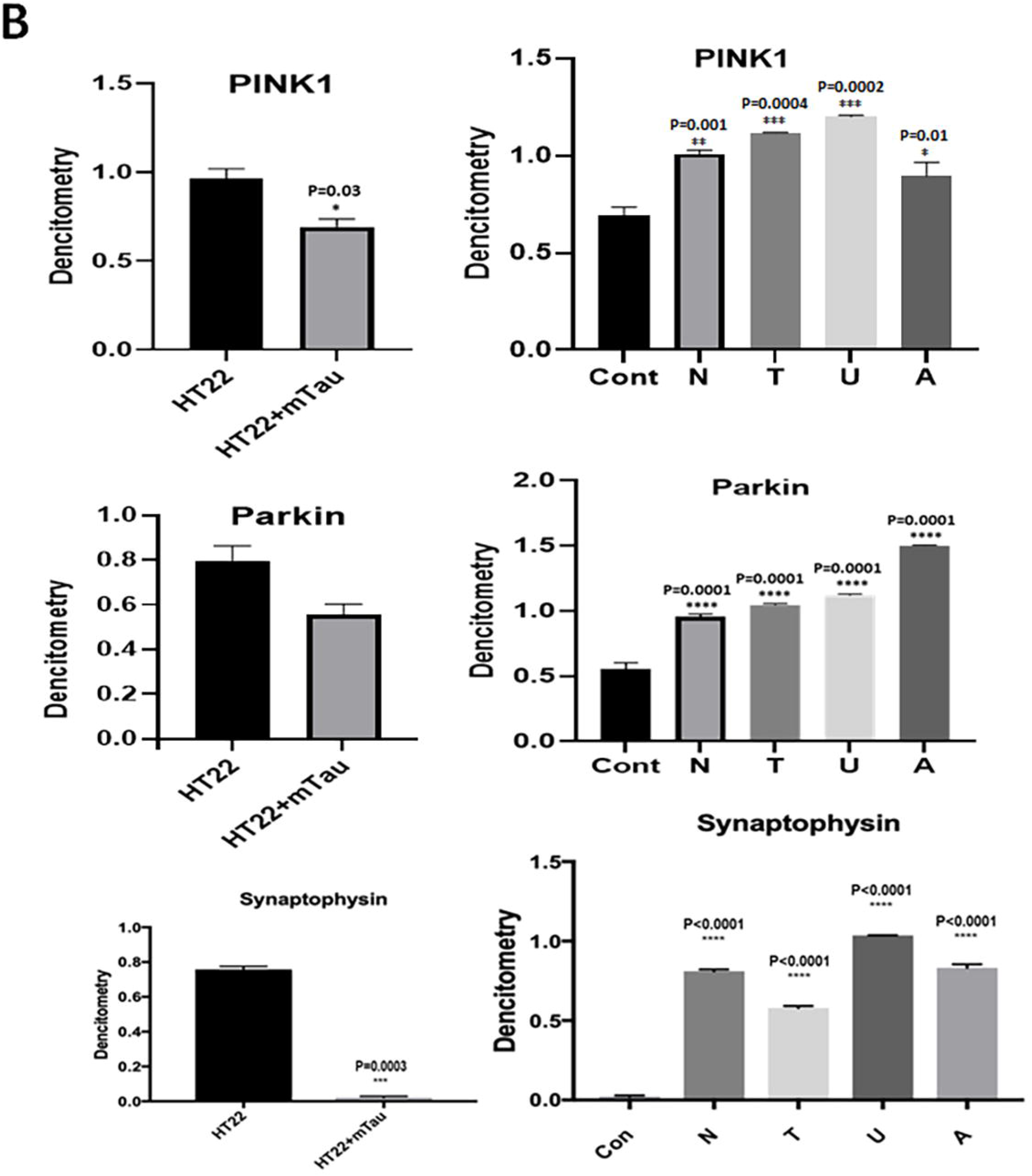
Western blot analysis of mitophagy and synaptic proteins. **(A)** Representative immunoblots for control HT22 cells and mTau-HT22 cells with or without mitophagy enhancers. **(B)** Represents quantitative densitometry analysis of mitophagy and synaptic proteins. Upon mTau transfection significant reduction were seen in the levels of PINK1, Parkin and synaptophysin (P=0.003) But levels of mitophagy and synaptic proteins increased with mitophagy enhancers treatment.

### Mitophagy enhancers increases synaptic protein synaptophysin

Our results show that expression of synaptic protein synaptophysin (P=0.0003) was significantly decreased in mTau-HT22 cells compared to control HT22 cells; however, synaptophysin was significantly increased in mitophagy enhancers treated mTau-HT22 cells compared to mitophagy enhancers untreated mutant tau cells. The relative P values as per the treatment is: Nicotinamide riboside P=0.0001, tomatidine P=0.0001; urolithin A P=0.0001, actinonin P=0.0001 (Figure 5A and 5B). Interestingly, statistical significance level was higher for urolithin A treated mTau-HT22 cells. Our results strongly suggest that mitophagy enhancers induces synaptic activity in the presence of mutant Tau, urolithin A is the best mitophagy enhancer out of others.

### Mitophagy enhancers increases cell survival

We performed the cell survival assay to see the effect of mitophagy enhancers on cell survival in HT22 cells and HT22 cells transfected with mutant tau cDNA. As presented in Figure 6 cell survival was significantly decreased in mTau-HT22 cells (P=0.0006) compared to control HT22 cells. However, cell survival was significantly increased in mitophagy enhancers treated mutant Tau-HT22 cells (nicotinamide riboside, P=0.0008, urolithin A, P=0.0001, tomatidine P=0.0001) as compared to mitophagy enhancers untreated mTau-HT22 cells.

**Figure 6.**
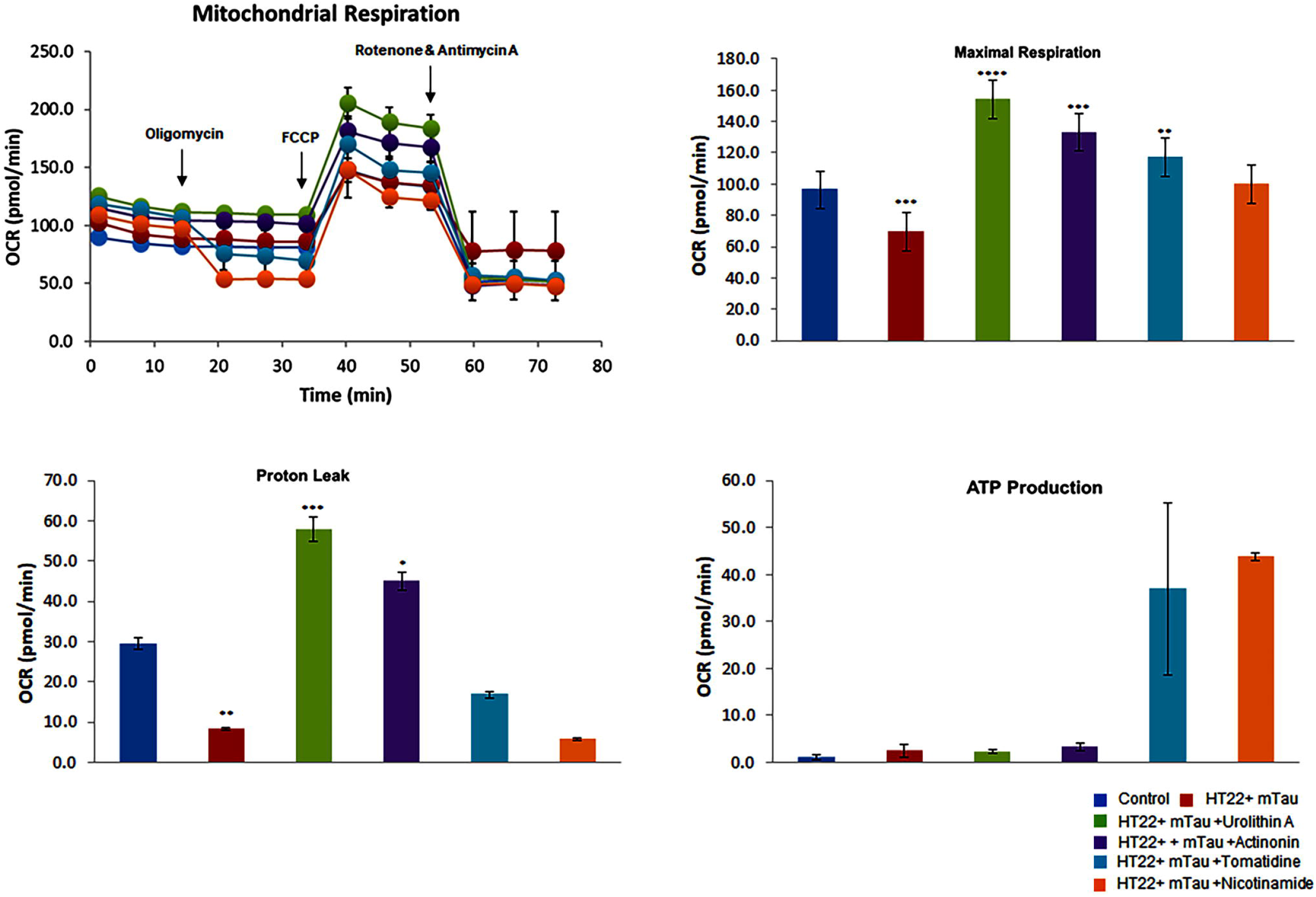
Cell survival assays in HT22 cells and HT22 cells transfected with mutant Tau cDNA. **(A)** Cell survival was significantly decreased in mTau-HT22 cells (P=0.0001) relative to control HT22 cells. However, cell survival was increased in mitophagy enhancers treated mutant Tau cells (nicotinamide riboside (P= 0.0001 and urolithin A (P=0.0001) relative to mitophagy enhancers untreated mutant Tau cells.

### Mitophagy enhancers increased maximal respiration in mutant Tau-HT22 cells

Additionally, we measured the mitochondrial respiration using Sea Horse Bioanalyzer in HT22 cells transfected with mutant Tau cDNA and treated with mitophagy enhancers. As shown in Figure 7, maximal OCR was decreased in mTau-HT22 cells (P=0.0006) compared to control, HT22 cells. However, maximal OCR was significantly increased in mTau-HT22 cells treated with mitophagy enhancers (urolithin A, P=0.0001; actinonin, P=0.0002; tomatidine, P=0.001) compared to untreated HT22 cells. Our results suggest that all mitophagy enhancers increases maximal OCR level.

**Figure 7:**
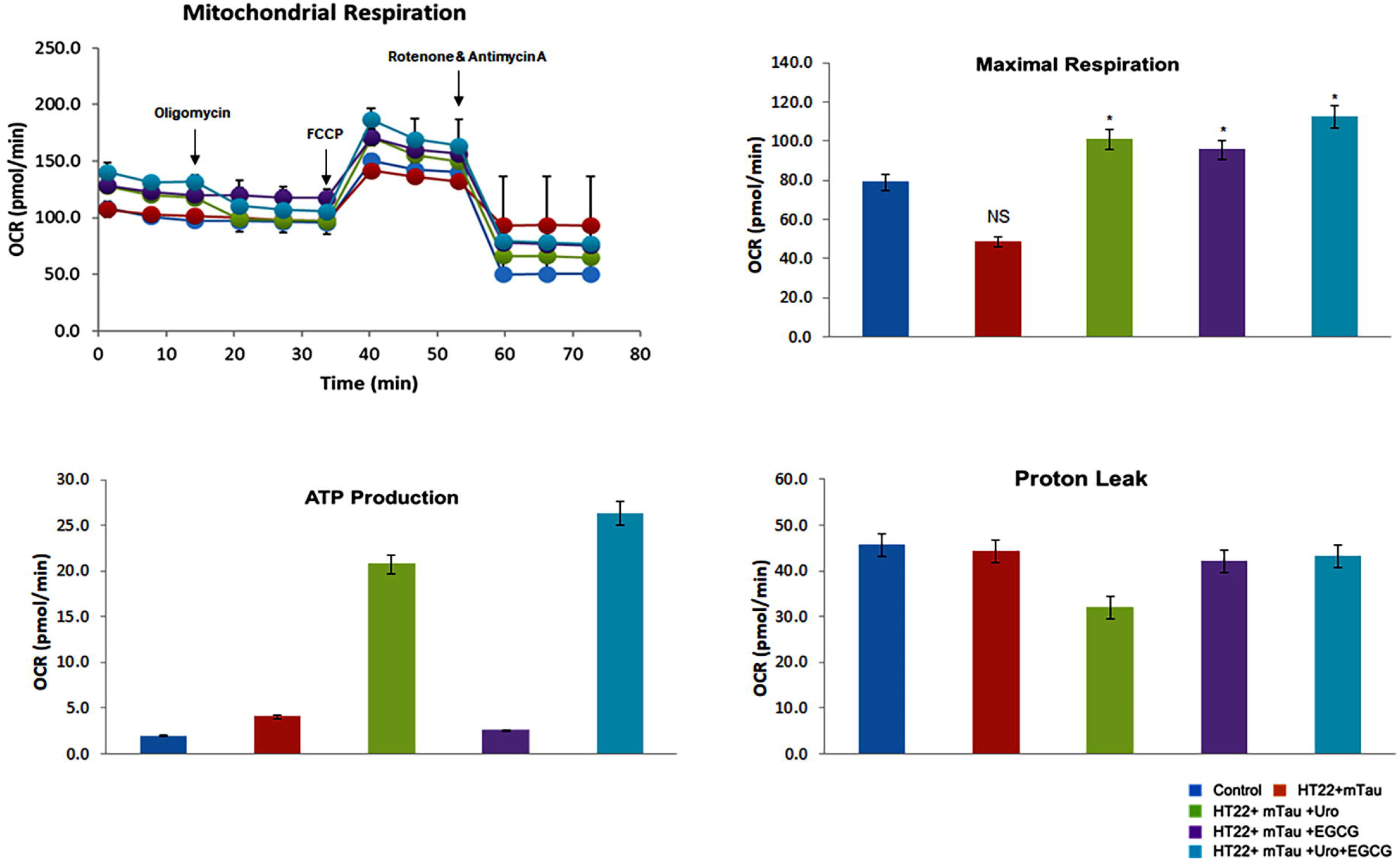
Mitochondrial respiration using mitophagy enhancers treated HT22 cells and mutant TauHT22 cells. The maximal oxygen consumption rate (OCR) was assessed in cells treated with mitophagy enhancers urolithin A (10µM), actionin (2µM), Tomatidine (1µM) and NAD (2µM) HT22 cells using an XFe96-well Extracellular Flux Analyzer (Seahorse Bioscience). Significantly increased maximal OCR was observed in HT22 cells treated with mitophagy enhancers relative to untreated HT22 cells, indicating that all mitophagy enhancers showed increased maximal OCR; however, urolithin A maximal respiration is the highest. The maximal (OCR, ATP and proton leaks were assessed in mTau-HT22 cells treated with mitophagy enhancers urolithin A (10µM), actinonin (2µM), tomatidine (1µM) and nicotinamide riboside (2µM) using Seahorse Bionalyzer. Increased maximal OCR was observed in mTau-HT22 cells treated with urolithinA, actionin, tomatidine and nicotinamide riboside relative to untreated mTauHT22 cells, indicating that mitophagy enhancers showed increased maximal OCR and ATP.

### Mitophagy enhancers increased maximal respiration in mutant tau-HT22 cells – Urolithin A, EGCG and Urolithin A + EGCG

To determine urolithin A, EGCG (green leaf extract) and a combination of both urolithin A + EGCG, we measured the mitochondrial respiration using Sea Horse Bioanalyzer in HT22 cells transfected with mutant Tau cDNA and treated with urolithin A, EGCG and a combination of both urolithin A + EGCG. As shown in Figure 8, maximal OCR was significantly increased in mtau-HT22 cells treated with urolithin A (P=0.02); EGCG (P=0.03) and a combination of urolithin A + EGCG (P=0.01 compared to untreated HT22 cells. As expected, a combination of urolithin A + EGCG treated mTau-HT22 cells showed a strongest protective effective, indicating combination therapy is powerful.

**Figure 8:**
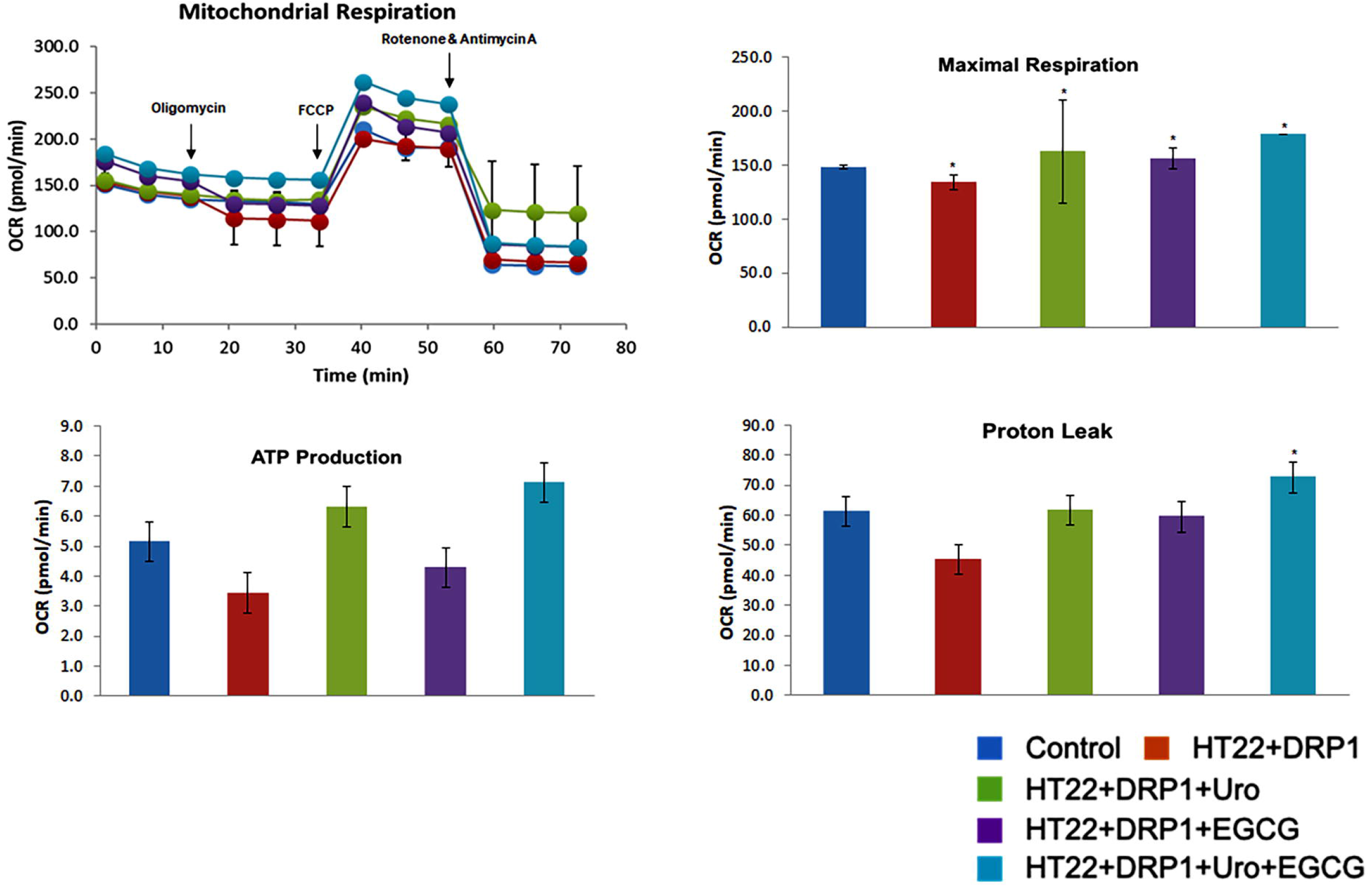
Urolithin A, EGCG and a combination of urolithin A+EGCG. The mitochondrial respiration was assessed using Sea Horse Bioanalyzer in HT22 cells transfected with mutant Tau cDNA and treated with urolithin A, EGCG and a combination of both urolithin A + EGCG. The maximal OCR was significantly increased in mTau-HT22 cells treated with urolithin A (P=0.02); EGCG (P=0.03) and a combination of urolithin A + EGCG (P=0.01) compared to untreated HT22 cells. As expected, a combination of urolithin A + EGCG treated mTau-HT22 cells showed a strongest protective effective, indicating combination therapy is powerful.

### Mitophagy enhancers increased maximal respiration in HT22 cells transfected with full-length Drp1 and treated with mitophagy enhancers

In our next experiment, we assessed mitophagy enhancers (urolithin A and EGCG) on mitochondrial respiration in HT22 cells transfected with full length Drp1 and treated with mitophagy enhancers using Sea Horse Bioanalyzer (Figure 9). The maximal OCR was decreased in HT22 cells transfected with full length Drp1 (P=0.01) relative to control, HT22 cells (Figure 9). On the other hand, maximal OCR was significantly increased in HT22 cells transfected with full length Drp1 and treated with mitophagy enhancers (urolithin A P=0.02; EGCG P=0.02; urolithin A+EGCG P=0.01) to untreated HT22 cells, indicating that all mitophagy enhancers showed enhanced maximal OCR.

**Figure 9:**
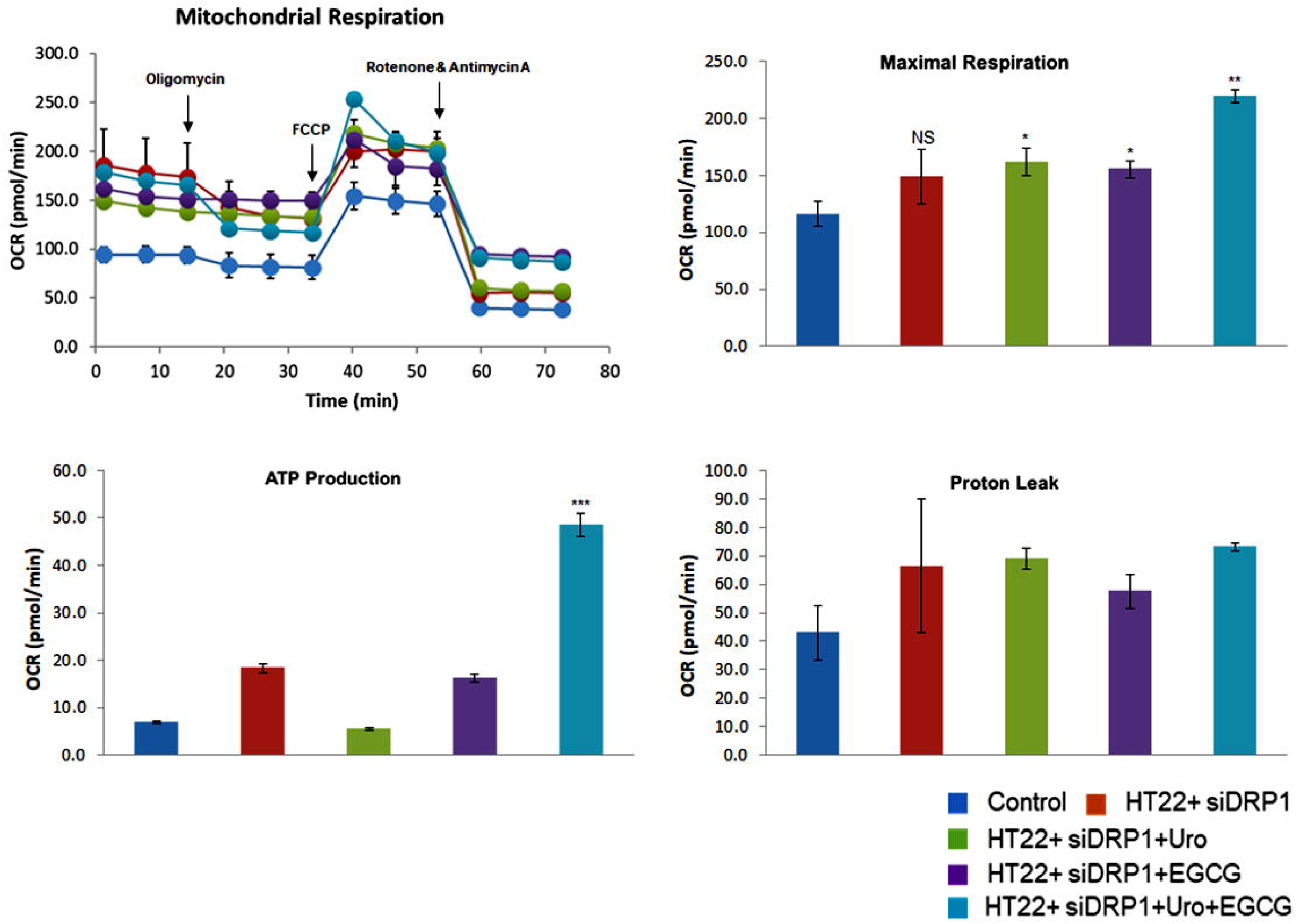
Mitochondrial respiration in full length Drp1 overexpressed and mitophagy enhancers treated HT22 cells. To determine the effects of mitophagy enhancers on mt respiration, Mito-stress assay was performed using an XFe96-well Extracellular Flux Analyzer (Seahorse Bioscience). The maximal oxygen consumption rate (OCR) was measured in full length Drp1 overexpressed and mitophagy enhancers (urolithin A and EGCG) HT22 cells. As shown in Figure 9, significantly decreased maximal OCR in full length Drp1 overexpressed HT22 cells. Increased maximal OCR in full length Drp1 overexpressed and treated with mitophagy enhancers HT22 cells, indicated that all mitophagy enhancers showed increased maximal OCR. However, combine effect of urolithin A and EGCG maximal respiration is the highest.

### Mitophagy enhancers increased maximal respiration in RNA silenced Drp1 and mitophagy enhancers treated HT22 cells

As shown in Figure 10, we assessed mitophagy enhancers (urolithin A and EGCG) on mitochondrial respiration in RNA silenced Drp1-HT22 cells and treated with mitophagy enhancers using Sea Horse Bioanalyzer. The maximal OCR was significantly increased in RNA silenced Drp1 and mitophagy enhancers treated HT22 cells (urolithin A P=0.03; EGCG P=0.02; urolithin A+EGCG P=0.001) to untreated HT22 cells (Figure 10), indicating that all mitophagy enhancers showed enhanced maximal OCR.

**Figure 10:**
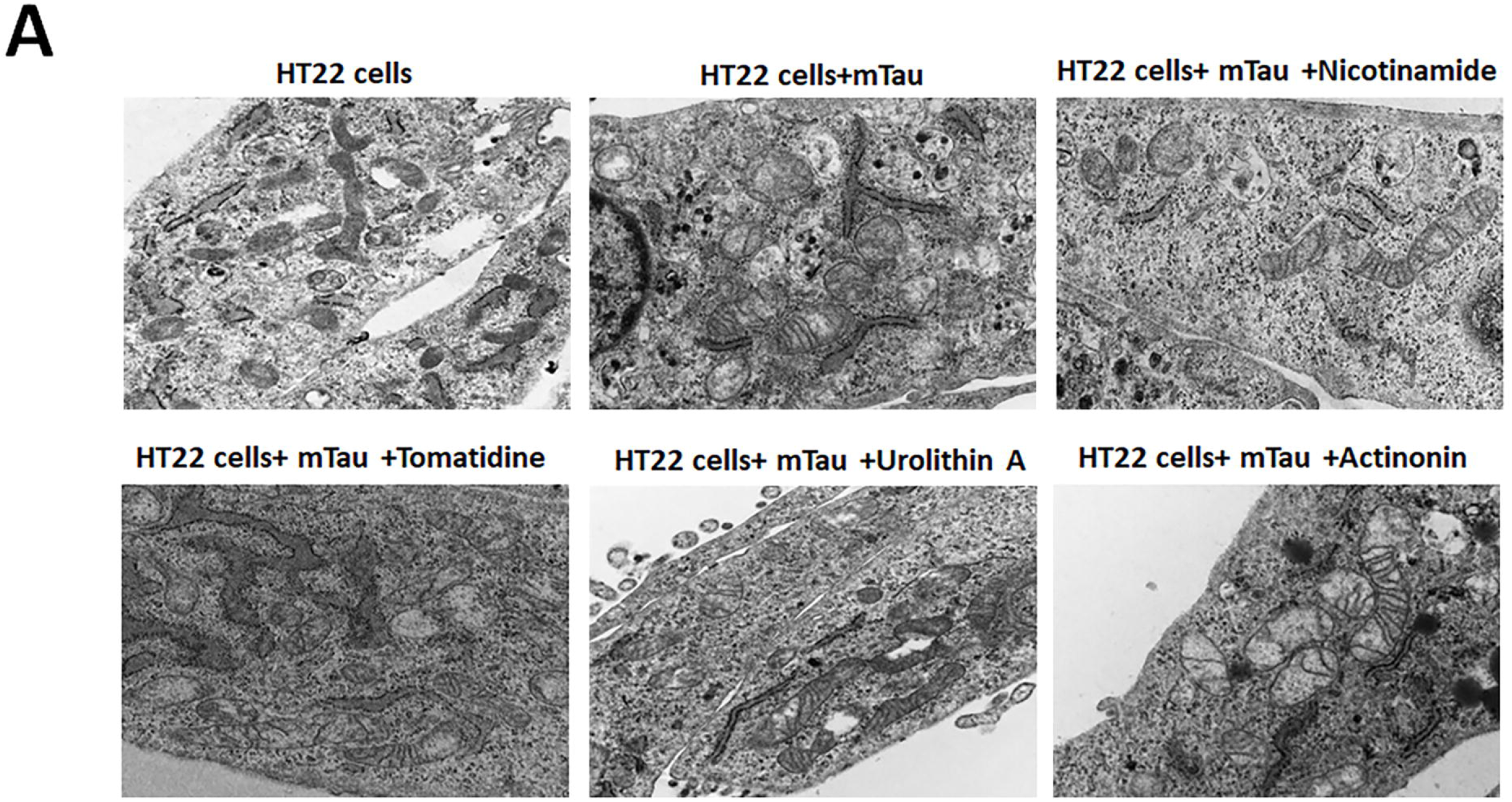
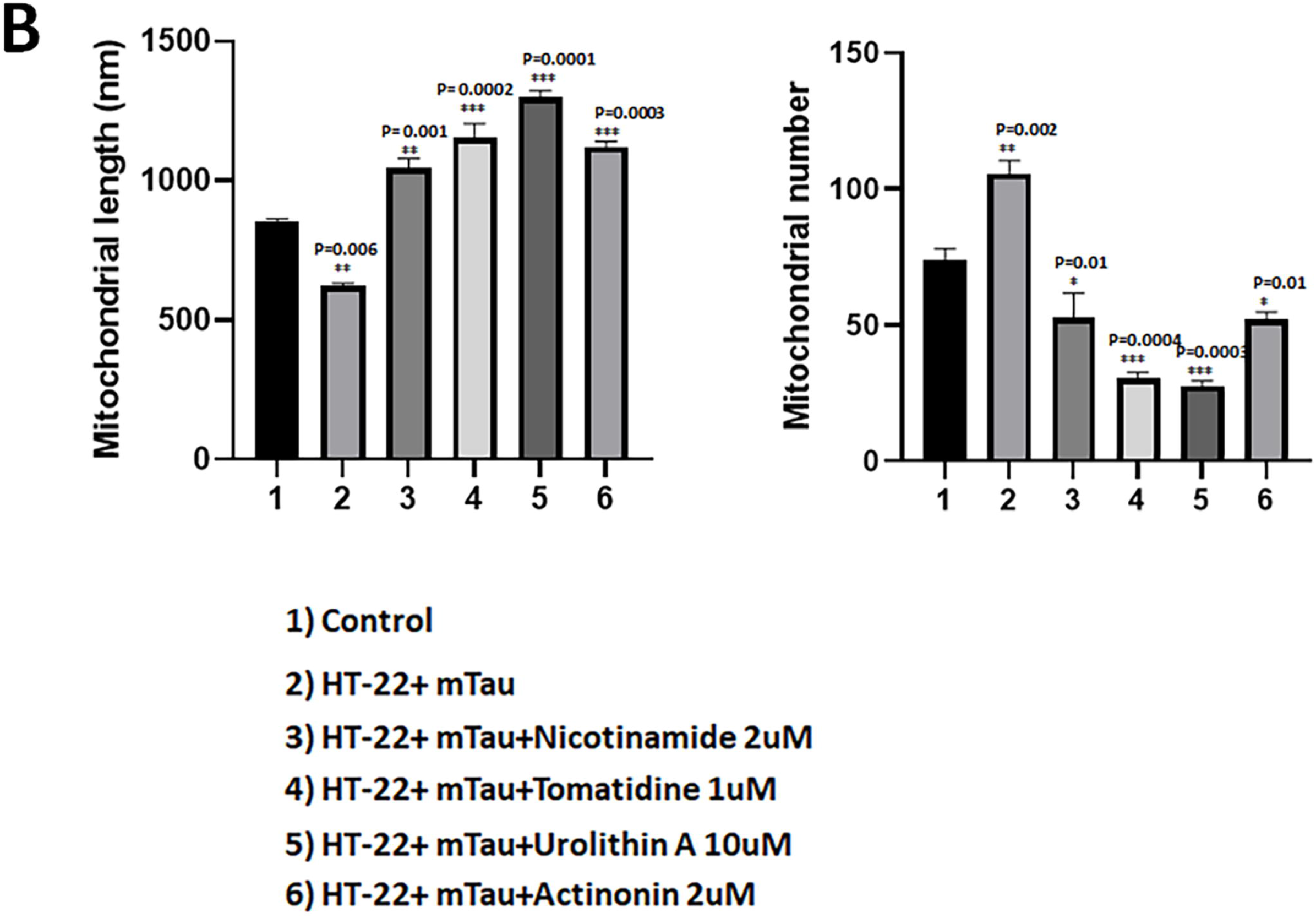
Mitochondrial respiration in RNA silenced Drp1 and mitophagy enhancers treated HT22 cells. Figure 10 shoes, increased maximal oxygen consumption rate (OCR) in Drp1 RNA silenced HT22 cells and treated with mitophagy enhancers (urolithin A and EGCG). Combination effect of (urolithin A + EGCG) maximal respiration is the highest as compared to single dose.

### Transmission electron microscopy analysis and mitochondrial length and number

Transmission electron microscopy technique was used to see the effects of mutant Tau on mitochondrial number and length; we compared mTau-HT22 cells and untransfected control-HT22 cells.

#### Mitochondrial number in mutant Tau-HT22 cells

Significant increased number of mitochondria (P=0.002) and reduced mitochondrial length (P=0.006) were observed in mTau-HT22 cells as compared to non-transfected, control-HT22 cells (Figure 11A and 11B), which suggests that tau fragments mitochondria.

**Figure 11:**
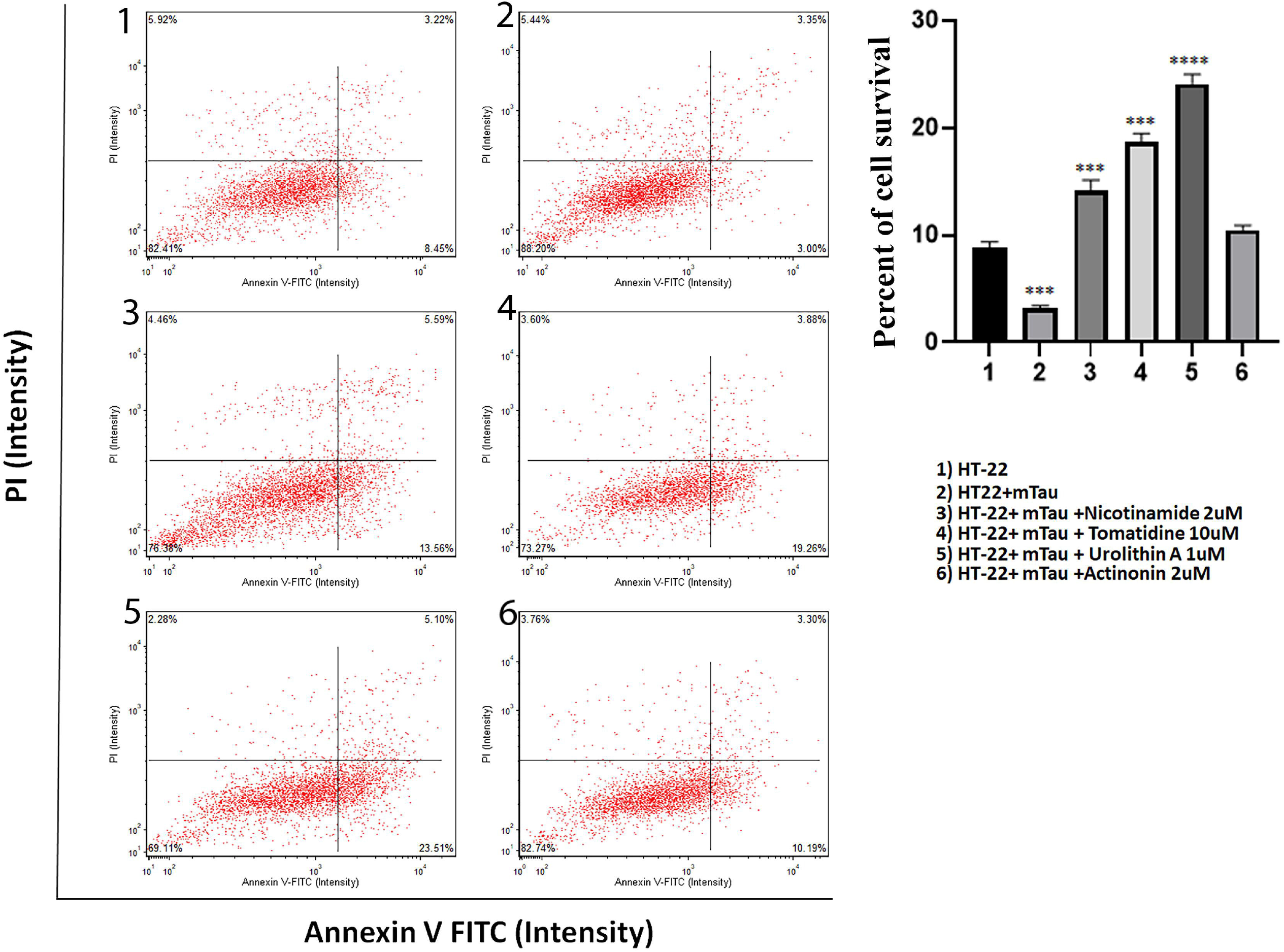
Transmission electron microscopy analysis. Mitochondrial number and length in control HT22 cells and mutant Tau cDNA transfected HT22 and treated with mitophagy enhancers for 24 hours. **(A)** Representative transmission electron microscopy images of mitochondria in the untreated HT22 cells and mitophagy enhancers treated mTau-HT22 cells **(B)** Quantitative analysis of mitochondrial number and length in each of the 6 groups. Significantly increased number of mitochondria were found in HT22 cells transfected with mutant Tau relative to untransfected HT22 cells. Mitochondrial length significantly decreased upon mutant Tau cDNA transfection. mitophagy enhancers treatment decreased the mitochondrial number and increased its length in the mTau-HT22 cells.

Additionally, we analyzed mitochondrial number and length in mTau-HT22 cells treated and untreated with mitophagy enhancers (urolithin A, actinonin, tomatidine and nicotinamide riboside). It is noticed that mitochondria number is significantly decreased in mTau-HT22 cells treated with nicotinamide riboside (P=0.01), tomatidine (P=0.0004), urolithin A (P=0.0003) and actinonin (P=0.01) compared to mitophagy enhancers untreated mtau-HT22 cells. Mitochondrial length is also significantly increased in mitophagy enhancers treated mTau-HT22 cells compared to mitophagy enhancers untreated mTau-HT22 cells; nicotinamide riboside (P=0.001), tomatidine (P=0.0002), urolithin A (P=0.0001), and actinonin (P=0.0003). Our findings suggest that mitophagy enhancers maintain and/or boost quality of mitochondria in AD neurons.

## Discussion

The long-term goal of our lab studies is to develop effective mitochondrial therapies for Alzheimer’s disease (AD), with a focus on mitochondria and synapse in relation to amyloid beta and phosphorylated tau. Two decades of research from our lab and others revealed that mitochondrial abnormalities, including defective mitochondrial dynamics, impaired mitochondrial biogenesis, defective axonal transport of mitochondria and impaired clearance of dead mitochondria are linked to phosphorylated and amyloid beta in AD^4-17, 26-37^. Increased proliferation of dead mitochondria at synapse reported to disrupt synaptic and cognitive functions AD patients. We and others proposed that increased clearance of dead mitochondria is an ideal therapeutic strategy for AD^8,9,43^. Mitophagy enhancers are potential candidates to improve mitochondrial and functions.

Our lab recently optimized the doses of mitophagy enhancers including urolithin A, actinonin, tomatidine, nicotinamide riboside and assessed mitophagy enhancers against amyloid beta-induced synaptic and mitochondrial studies^40^. Interestingly, findings suggest that mitophagy enhancers reduce the mitochondrial and synaptic toxicities and improve bioenergetics and cell survival of amyloid beta expressed neurons ^40^.

In the current study, we treated mutant Tau expressed in HT22 (mTau-HT22) cells with mitophagy enhancers and assessed cell survival, mitochondrial bioenergetics/respiration, mRNA and protein levels of mitochondrial/synaptic genes. We also assessed mitochondrial morphology in mTau-HT22 cells. Mutant Tau-HT22 cells treated with mitophagy enhancers increased mitochondrial respiration, cell survival and also increased fusion, synaptic & mitophagy activities. Further, urolithin A showed strongest protective effects among all enhancers against mTau-induced mitochondrial and synaptic toxicities in AD. In addition, our combination treatments of urolithin A + EGCG (green leaf extract), revealed significantly increased mitochondrial respiration against mutant Tau or phosphorylated tau toxicities.

### Normal Tau and phosphorylated Tau in AD

Normal tau is critical for microtubule binding, axonal transport of organelles from soma to nerve terminals and neuronal sprouting and synaptic and functions^20^. Although genetic mutations are not directly involved with AD development, in AD state tau becomes hyperphosphorylated, not able bind microtubules, leading to impaired axonal transport and synaptic starvation, leading to cognitive decline^20^. In addition, our lab reported phosphorylated tau physically interacts with mitochondrial fission protein Drp1, induces GTPase Drp1 enzymatic activity and causes excessive mitochondrial fragmentation and defective mitophagy ^9,26^. In addition, phosphorylated tau also interacted with mitochondrial outermembrane protein, VDAC1 and causes impairment(s) in mitochondrial gating, ultimately leading to mitochondrial dysfunction and mitophagy defects in AD neurons^28^. Therefore, the use of mitophagy enhancers in the current is highly relevant.

### Protective effects of mitophagy enhancers

In the current study, we assessed mRNA and protein levels of mitochondrial dynamics, biogenesis, synaptic and mitophagy genes in mTau-HT22 cells treated and untreated with enhancers. Regarding mitochondrial dynamics, mRNA levels of mitochondrial fission genes were significantly increased and fusion genes were reduced in the mTau-HT22 cells. These were reverse in all mitophagy enhancers treated cells. However, urolithin A treated cells showed higher fold change differences. These results indicate that urolithin A maintain mitochondrial dynamics in a best possible way among all mitophagy enhancers tested drugs in our study.

Regarding, mitochondrial biogenesis, mTau-HT22 cells showed reduced mitochondrial biogenesis relative control HT22 cells, indicating that mTau affects mitochondrial biogenesis: On the other hand, in mitophagy enhancers treated mTau-HT22 cells showed increased mitochondrial biogenesis activity. Urolithin A enhanced biogenesis activity higher than all other enhancers studied.

PINK 1 and Parkin proteins are important to maintain mitophagy in both healthy and disease states, In AD, PINK 1 and Parkin levels are low, leading impaired clearance of dead mitochondria, therefore, it is important to enhance and/or at least maintain these proteins ^8,9,43^. As we observed, mRNA and protein levels of PINK1 and Parkin reduced in mTau-HT22, indicating mutant Tau reduces these proteins. However, these proteins were increased mitophagy enhancers treated mTau-HT22 cells. Similar to mitochondrial biogenesis, urolithin A showed strongest protective effect even for mitochondrial dynamics in mTau-HT22 cells.

Synaptic protein, synaptophysin was reduced in mTau-HT22 cells relative to control HT22 cells, indicating that mutant Tau effects synaptic activity, it has been extensively reported in AD ^45,46^. However, synaptophysin levels were increased in mitophagy enhancers treated mTau-HT22. Urolithin A treated cells showed strongest effect for synaptophysin among all mitophagy enhancers.

### Mitophagy enhancers reduce full-length mutant tau

In the current study, we found, a full-length 70-kDa pTau protein in mutant Tau transfected HT22 cells. On the other hand, full-length mutant Tau levels were reduced in the mitophagy enhancers treated cells and urolithin A showed highest decrease of mutant full-length mutant Tau, representing that urolithin A diminishes full-length mutant Tau.

### Mitophagy enhancers increases cell survival

Cell survival was significantly reduced in mTau-HT22 cells relative to control cells, and this was reversed in mitophagy enhancers treated cells. Cell survival was increased with a much stronger in urolithin A treated mTau-HT22 cells. These observations indicate that mitophagy enhancers are good candidates and urolithin A is the best among all we studied.

### Increased mitochondrial respiration in mutant Tau-HT22 cells treated with mitophagy enhancers

In the current study, we extensively assessed mitochondrial respiration using Sea Horse Bioanalyzer in mTau-HT22 cells. As expected mTau-HT22 cells showed reduced mitochondrial respiration. On the other hand, mitochondrial respiration (maximal OCR, ATP) was increased in mitophagy enhancers treated mTau-HT22 cells and urolithin A showed the strongest effect.

We also assessed, if combined therapy has any effect on mitochondrial respiration, in order to understand this, we treated mTau-HT22 cells with 1) urolithin A, 2) EGCG (green leaf extract) and 3) combination of both urolithin A and EGCG. Maximal OCR was significantly increased in mTau-HT22 cells treated with urolithin A, EGCG and a combination of urolithin A + EGCG compared to untreated HT22 cells. As expected, a combination of urolithin A + EGCG treated mTau-HT22 cells showed a strongest protective effective, indicating combination therapy is powerful.

### Drp1 overexpression and RNAsi effects on mitochondrial respiration

To determine the role of mitochondrial fission protein, Drp1 on mitochondrial respiration in mTau-HT22 cells and mitophagy enhancers, urolithin A, EGCG and in combination, we assessed mitochondrial respiration in HT22 cells transfected with full length mouse Drp1 cDNA. The maximal OCR was decreased in HT22 cells transfected with full length Drp1 relative to control cells. On the other hand, maximal OCR was significantly increased in HT22 cells transfected with full length Drp1 and treated with mitophagy enhancers, urolithin A, EGCG and a combination of urolithin A+EGCG and the effect was stronger in a combination treatment.

We also assessed mitophagy enhancers (urolithin A, EGCG and combination) on mitochondrial respiration in RNA silenced mouse Drp1-HT22 cells and treated with mitophagy enhancers using Sea Horse Bioanalyzer. The maximal OCR was significantly increased in RNA silenced Drp1 and mitophagy enhancers treated HT22 cells to untreated HT22 cells. Overall, these data suggest that RNA silenced Drp1 enhances mitochondrial respiration, on the other hand Drp1 overexpressed cells showed reduced mitochondrial respiration, in other words increased Drp1 is toxic to cells.

Our extensive transmission electron microscopy analysis data indicate mitophagy enhancers reduced fragmented mitochondria and increased mitochondrial length, this effect is stronger in urolithin A treated mTau-HT22 cells.

## Authors Contributions

S.S. and P.H.R contributed to the conceptualization and formatting of the article. S.S. H.M. and N. S. conducted experiments and S.S. A.P.R and P.H.R are responsible for writing, original draft preparation, and finalization of the manuscript. A.P.R and P.H.R. is responsible for funding acquisition.

## Availability of data and materials

Not applicable

## Compliance to ethical standards

Presented research is compliance with ethical standards

## Ethics Approval

Not applicable

## Consent to participate

Not applicable

## Consent to publication

All authors agreed to publish the contents

